# *Mycena* genomes resolve the evolution of fungal bioluminescence

**DOI:** 10.1101/2020.05.06.079921

**Authors:** Huei-Mien Ke, Hsin-Han Lee, Chan-Yi Ivy Lin, Yu-Ching Liu, Min R. Lu, Jo-Wei Allison Hsieh, Chiung-Chih Chang, Pei-Hsuan Wu, Meiyeh Jade Lu, Jeng-Yi Li, Gaus Shang, Rita Jui-Hsien Lu, László G. Nagy, Pao-Yang Chen, Hsiao-Wei Kao, Isheng Jason Tsai

## Abstract

Mushroom-forming fungi in the order Agaricales represent an independent origin of bioluminescence in the tree of life, yet the diversity, evolutionary history, and timing of the origin of fungal luciferases remain elusive. We sequenced the genomes and transcriptomes of five bonnet mushroom species (*Mycena* spp.), a diverse lineage comprising the majority of bioluminescent fungi. Two species with haploid genome assemblies ∼150Mb are amongst the largest in Agaricales, and we found that a variety of repeats between *Mycena* species were differentially mediated by DNA methylation. We show that bioluminescence evolved in the last common ancestor of mycenoid and the marasmioid clade of Agaricales and was maintained through at least 160 million years of evolution. Analyses of synteny across genomes of bioluminescent species resolved how the luciferase cluster was derived by duplication and translocation, frequently rearranged and lost in most *Mycena* species, but conserved in the *Armillaria* lineage. Luciferase cluster members were co-expressed across developmental stages, with highest expression in fruiting body caps and stipes, suggesting fruiting-related adaptive functions. Our results contribute to understanding a *de novo* origin of bioluminescence and the corresponding gene cluster in a diverse group of enigmatic fungal species.

**Significance:** We present the genomes of five new bonnet mushroom *Mycena* species, formerly the last fungal bioluminescent lineage lacking reference genomes. These genome-scale datasets allowed us to construct an evolutionary model pinpointing all possible changes in the luciferase cluster across all fungi and additional genes involved in bioluminescence. We show that luciferase clusters were differentially lost in different fungal lineages and in particular a substantial loss was observed in the *Mycena* lineage. This can be attributed to genome regions of *Mycena* underwent different evolutionary dynamics. Our findings offer insights into the evolution of how a gene cluster that emerged 160 million years ago and was frequently lost or maintained due to differences in genome plasticity.

## Introduction

The genus *Mycena* (Pers.) Roussel, comprises approximately 600 small mushroom species widely distributed around the world(1). Also known as bonnet mushrooms, *Mycena* species are usually characterised by a bell-shaped cap, a thin stem (**Fig. 1A**), and a gilled or porioid hymenophore(2). *Mycena* also have a diversity of life history strategies; while many species are saprotrophic, they can be pathogens as well as mycorrhizal(3). Despite its vast diversity of lifestyles and phenotypes, there are many questions concerning the basic biology, ecology and genomics of this genus. One particular fascinating trait is bioluminescence, which is most widespread in the genus *Mycena*. At least 68 of the 81 known bioluminescent fungi belong to *Mycena* (4), yet these account for less than 12% of the ca. 600 *Mycena* species (5), suggesting an intricate loss/gain history and potential convergence within the genus.

**Fig. 1.**
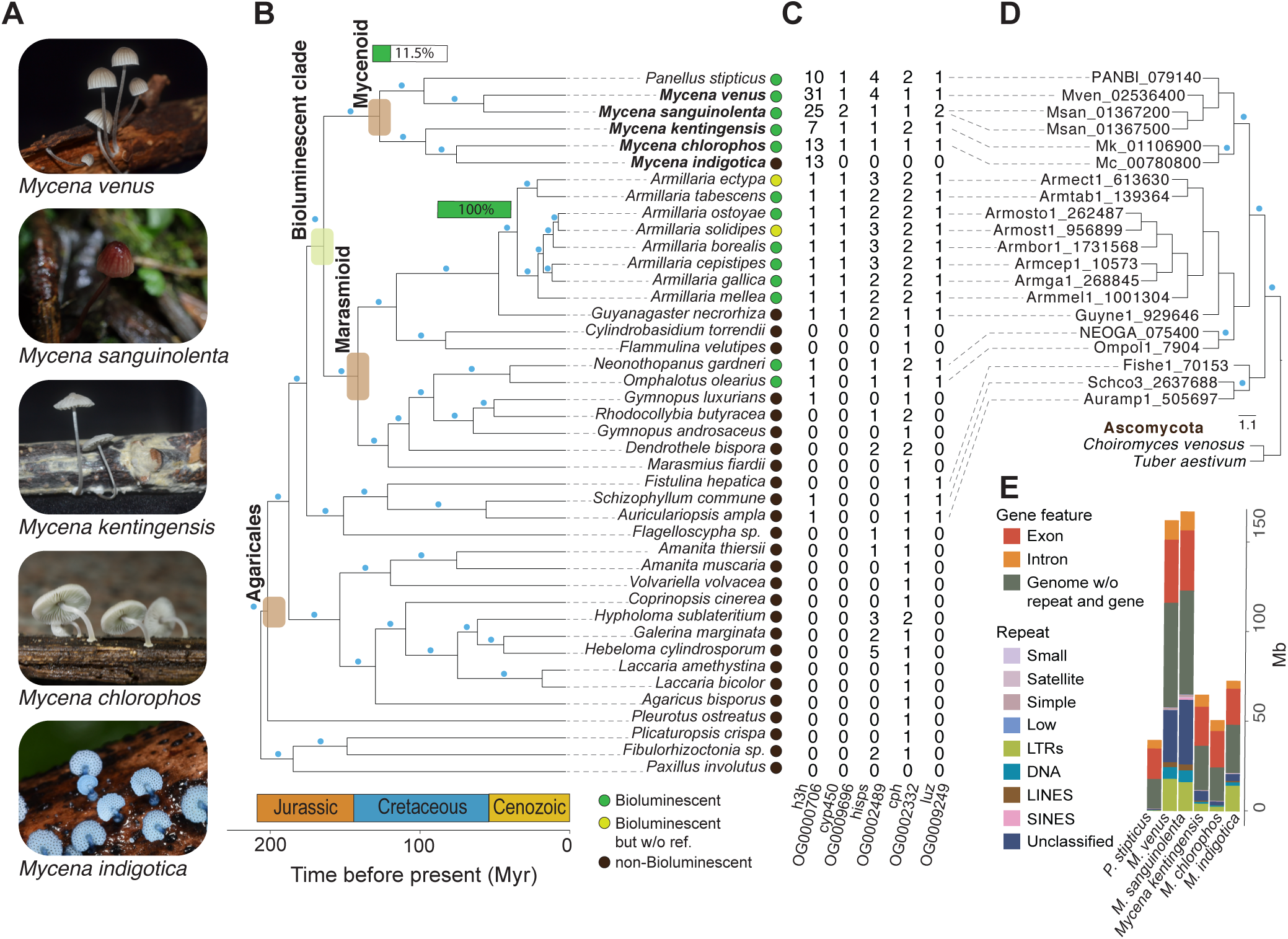
Phylogenomic analysis of *Mycena* and related fungi. (*A*) The five species sequenced in this study. (*B*) Species trees inferred from a concatenated supermatrix of the gene alignments using the 360 single-copy orthogroups. X-axis denotes divergence time estimates. Blue dot on a branch indicates a bootstrap value > 90. Green horizontal bars indicate the percentage of bioluminescent fungi found in either the mycenoid or the *Armillaria* lineage. (*C*) Gene copy number in the orthologous groups (OG) associated with luciferin biosynthesis pathway including luciferase (*luz*), hispidin-3-hydroxylase (*h3h*), hispidin synthase (*hisps*), cytochrome P450 (*cyp450*) and caffeylpyruvate hydrolase (*cph*). (*D*) Reconciliated phylogeny of fungal luciferase. Blue dot on a branch indicates a bootstrap value > 90. (*E*) Haploid genome sizes for 42 species broken down by repeat types and gene features. Averaged content in the genomes of 14 outgroup species are indicated as one bar. Repeats including transposable elements (TEs): long terminal repeats (LTRs), long interspersed nuclear elements (LINES), short interspersed nuclear elements (SINEs), DNA transposons (DNA), and other types of repeats: small RNA (Small), simple repeats (Simple), and low complexity repeats (Low).

Fungal light emission involves two main steps. First, a luciferin precursor of hispidin is hydroxylated by hispidin-3-hydroxylase (H3H) into 3-hydroxyhispidin (luciferin) (6). Oxygen is then added to the luciferin by luciferase, producing a high energy intermediate which is subsequently decomposed, yielding light emission. Previously, Kotlobady *et al*. have identified the fungal luciferase, which is physically adjacent to these enzymes and forms a gene cluster containing luciferase, hispidin synthase and H3H (7). This cluster was found to be conserved across bioluminescent fungi of three lineages: *Armillaria*, mycenoid and *Omphalotus* (7). Phylogeny reconstruction suggested that luciferase originated in early Agaricales. *Armillaria* and *Omphalotus* belong to the marasmioid clade, whereas *Mycena* was recently found to be sister of the marasmioid clade(8). Recent genome sequencing efforts in the marasmioid clade revealed diverse genomic and life history traits, including genome expansion and pathogenicity in *Armillaria* spp.(9), novel wood decay strategies(10) or fruiting body development(11). Genomes of two *Mycena* species were sequenced(7), however, the fragmented assemblies (N50 5.8–16.7 kb) impeded comparative genomic analyses of features such as synteny(12). These resources provide a substrate for studies of genome evolution and of bioluminescence in fungi, however, several key questions remain unresolved. Here, we set out to understand how the luciferase cluster originated and was lost or retained, determine levels of variation in this cluster across these lineages, and identify novel genes involved in bioluminescence.

To gain insights into the evolution of fungal bioluminescence and the ecology of mycenoid species, we sequenced the genomes of four bioluminescent (*Mycena chlorophos, M. kentingensis, M. sanguinolenta* and *M. venus*) and one non-bioluminescent (*M. indigotica*) species. We conducted comparative genomics with representative genomes of all bioluminescent fungal clades, putting particular emphasis on genome-wide synteny to investigate the evolutionary dynamics of the luciferase gene cluster through hundreds of millions of years. The variability in genome sizes among *Mycena* is likely associated with the differential expansion of repeats in the genomes, potentially due to the differential control on repeat activity by DNA methylation. The transcriptome of bioluminescent mycelium contained the luciferase cluster and co-expression analyses identified specific genes likely to associate with the bioluminescence and development in *Mycena*. Based on comparative analyses from fifteen available genomes of bioluminescent fungi, we reconstructed and formulated a model for the evolution of fungal bioluminescence.

## Results

### Assemblies and annotations of five *Mycena* species

We sequenced the genomes of the bioluminescent fungi *Mycena chlorophos, M. kentingensis, M. sanguinolenta* and *M. venus*, as well as the non-bioluminescent *M. indigotica* (**Fig. 1A**). These species were chosen for their phylogenetic positions (*SI Appendix*, Fig. S1) and because they displayed different bioluminescence intensities. An initial assembly of each species produced from Oxford Nanopore reads of each species (*SI Appendix*, Table S1) using the Canu (13) assembler. Only *M. indigotica* was successfully isolated from a basidiospore, and the four species that were isolated from heterokaryotic mycelium yielded assemblies 1.3–1.8 times larger than haploid genome sizes estimated from Illumina reads using GenomeScope (14) (*SI Appendix*, Table S2). Mitochondrial genomes in these species were separately assembled into single circular contigs of 88.3–133 kb long (*SI Appendix*, Fig. S2), and haploid nuclear genomes were constructed. The assemblies were further polished using Illumina reads and had a consensus quality value (QV) of 31.1-36.8 (*SI Appendix*, Table S2), which is similar to the QVs of recently published nanopore assemblies in human (also polished with Illumina reads; 23.7–43.5)(15). These haploid nuclear genomes were 50.9–167.2 Mb long, and two of them were amongst the largest in Agaricales reported to date. The assemblies consisted of 30–155 contigs with N50 4.1–17.8 Mb (*SI Appendix*, Table S2), which were comparable to representative fungal reference assemblies (*SI Appendix*, Fig. S3) and allowed for synteny comparisons (12). Stretches of TTAGGG hexamers were identified at the end of scaffolds, indicating telomeric repeats commonly found in Agaricales (16, 17). The largest scaffolds in *M. indigotica* and *M. kentingensis* were telomere-to-telomere, indicating gapless chromosomes.

Using a combination of reference fungal protein homology support and mycelium transcriptome sequencing (Dataset S1), 13,940–26,334 protein encoding genes were predicted in the *Mycena* genomes using MAKER2(18) pipeline, and were 92.1–95.3% complete (*SI Appendix*, Table S3) based on BUSCO(19) analysis. Orthology inference using Orthofinder (20, 21) placed these genes models and those of 37 other Basidiomycota genomes (*SI Appendix*, Table S4) into 22,244 orthologous groups (OGs; *SI Appendix*, Table S5). Of these OGs, 44.3% contained at least one orthologue from another basidiomycete, while 15–29% of the proteomes in each *Mycena* species were species-specific (Dataset S2). The genome sizes were positively correlated with proteome sizes, with the largest (*M. sanguinolenta*) and smallest (*M. chlorophos*) varying two- and three-fold, respectively. Interestingly, the mitochondrial genomes were larger in species with smaller genomes, and this was because nine out of 16 genes had gained many introns (*SI Appendix*, Table S6 and Fig. S2).

### Interplay between transposable elements and DNA methylation in *Mycena*

Similar to other fungal genomes(22, 23), much of the variation in the *Mycena* nuclear genome sizes can be explained by repetitive DNA content (*SI Appendix*, Table S7). Only 11.7% of the smallest genome (*M. chlorophos*) was repeats, which is in stark contrast to the 39.0% and 35.7% in *M. sanguinolenta* and *M. venus*, respectively. The majority of transposable elements in *Mycena* were long terminal repeats (LTRs) retrotransposons (60–85%), followed by DNA transposable elements (11%–24%) (**Fig. 1E** and *SI Appendix*, Table S7). Interestingly, the larger genomes of *M. sanguinolenta* and *M. venus* contained the lowest proportion of LTRs (24.9 and 31.1%, respectively), but highest proportion of unclassified repeats (55.4 and 50.3%, respectively) (*SI Appendix*, Table S7). 16.6–36.5% of the unclassified repeat families shared 53.8–60.5% nucleotide identity with known transposable elements, suggesting they were degenerated copies which we defined as relic TEs (*SI Appendix*, Table S8). **Fig. 2A** shows that the largest assembled chromosome of *M. indigotica* exhibits high protein-coding gene content and low transposable element density at scaffold centres, which is typical of fungal chromosomes(24, 25). Such observations were consistent across large *Mycena* scaffolds (typically >1 Mb), suggesting that our assemblies were robust enough to capture evolutionary dynamics across chromosomes.

**Fig. 2.**
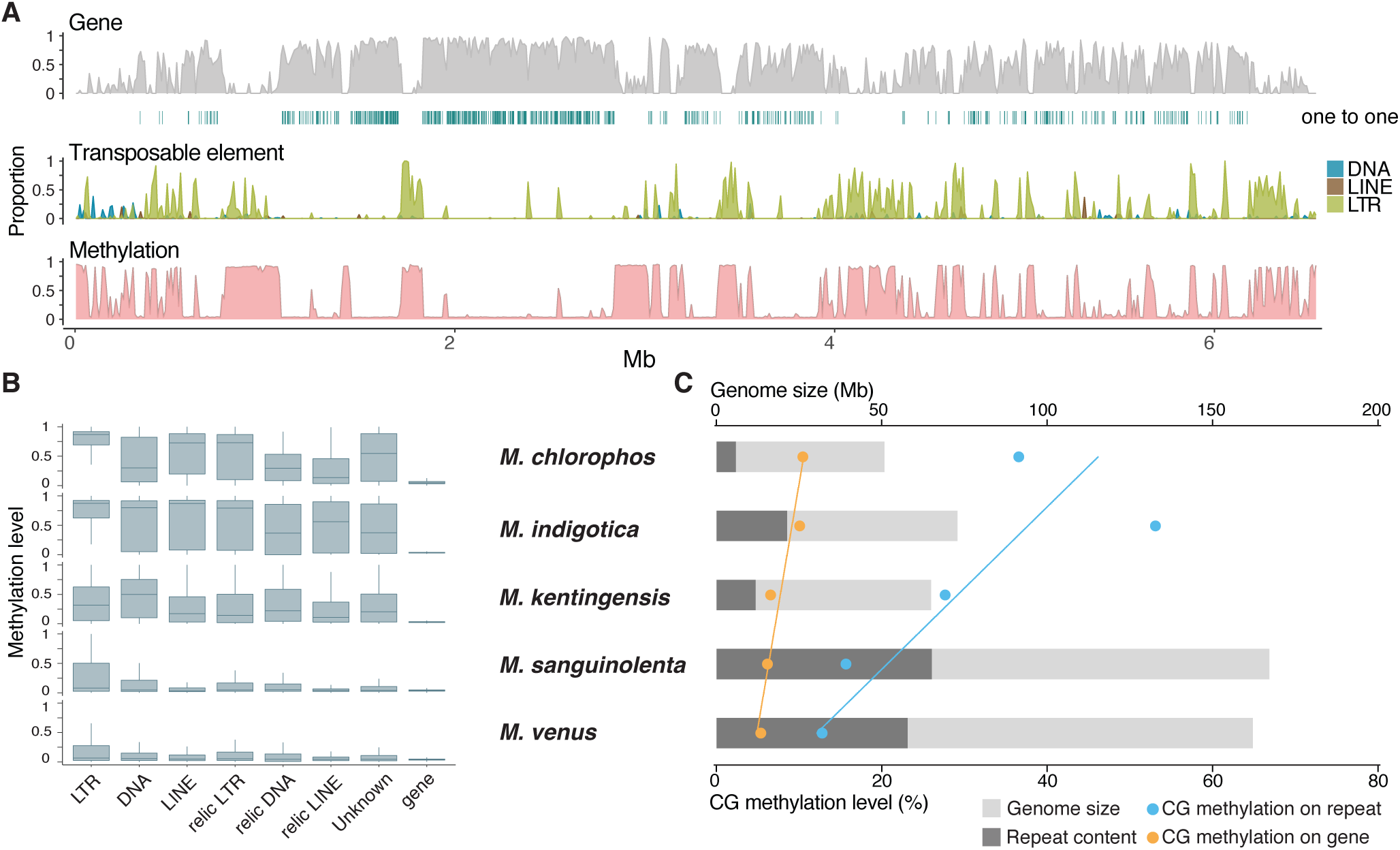
Distribution of *Mycena* genome features. (*A*) *M. indigotica* chromosome one. For every non-overlapping 10-kb window, the distributions from top to bottom are: (1) Gene density (percentage of nucleotides coverage). Green stripes denote positions of single-copy orthologue with *M. chlorophos*. (2) Density of transposable elements (TEs), including LTRs, LINES, and DNA. (3) Average methylation level called from CpG sites per window. The high methylation window generally clustered in high TE regions with low gene density. (*B*) The methylation level in genes and different types of repeats. (*C*) The relationships among genome size, number of repeats and CG methylation levels in *Mycena*.

We detected 5-methylcytosine (5mC) DNA methylation levels across the five *Mycena* assemblies with Nanopore long reads using deepsignal (26) which was initially trained with *M. kentingensis* bisulphite sequences (*Methods*). CG sites were found either highly (mCG level >60%) or weakly methylated (<15%) in gene body, displaying a bimodal distribution (*SI Appendix*, Fig. S4). Such a bimodal distribution has also been observed in plants, animals, and other fungi, including *Tuber melanosporum* and *Pseudogymnoascus destructans* (27-32). Within *Mycena*, the CG methylation in genes (5.4–10.5%) was much lower than that in repeats—i.e., TEs and unclassified repeats (11.6–84.5%) (**Fig. 2B**; *SI Appendix*, Table S9). The level of CG methylation in these genomes is comparable with those of a previous survey on DNA methylation in 528 fungal species (32), which revealed that 5mC levels were highest in Basidiomycota, further indicating that DNA methylation have a specific effect on repeats in *Mycena* genomes. DNA transposons or LTR were enriched in 5mC levels and were higher than flanking regions (*SI Appendix*, Fig. S5). Except for DNA transposons in *M. kentingensis*, LTR retrotransposons had the highest CG methylation levels of all types of transposable elements (**Fig. 2B**). Furthermore, CG methylation in relic TEs was clearly lower than that in classic TEs (*SI Appendix*, Table S9). Among the *Mycena* species, we found that *M. sanguinolenta* and *M. venus* with larger genomes and higher repeat content had lower levels of methylation in the repeats, and the repeat methylation was much higher in *M. indigotica, M. chlorophos*, and *M. kentingensis*, which have smaller genomes (**Fig. 2C**). The same pattern was also observed in genes, though they had fewer changes in their methylation level than did repeats. Our results indicate that the variant composition of repeats is differentially mediated by DNA methylation among closely-related *Mycena* species. Hence, genome expansion in *Mycena* was likely a result associated with transposable element proliferation and the accumulation of relic TEs, which yielded reduced methylation in active copies; this is also observed in some plants, e.g., *Arabis alpine*(33) and *Manihot esculenta* (34).

### A single origin of bioluminescent fungi in the ancestor of *Mycena* and the marasmioid clade

Phylogenomic analyses based on single-copy orthologue sets have placed *Mycena* sister to the marasmioid clade, including *Armillaria* and *Omphalotus*, which are the other two lineages in which bioluminescent species have been identified. This species phylogeny was recovered in both maximum likelihood analysis(35) of a concatenated supermatrix of single-copy gene alignments (**Fig. 1B**) and coalescent-based analysis using 360 gene trees(36) (*SI Appendix*, Fig. S6). In our four bioluminescent *Mycena* species, we identified genes involved in luciferin biosynthesis and their orthologues across species (**Fig. 1C**). **Fig. 1D** shows phylogenetic reconciliation, which suggests that the orthogroup containing luciferases was already present in the last common ancestor of the mycenoid+marasmioid clade and Schizophyllaceae, predating their incorporation into the luciferase cluster. This is in contrast to a previous report (7) suggesting that luciferase originated in the last common ancestor of the Agaricales. Phylogenies of other members of the luciferase cluster were also congruent with the species tree (*SI Appendix*, Fig. S7 A–D). Using MCMCtree (37) with three fossil calibrations, we estimated the age of mycenoid most recent common ancestor to be 105–147 million years ago (Mya) in the Cretaceous (**Fig. 1B**). This is consistent with recent estimates (78–110 (8) and mean 125 (1) Mya) and overlaps with the initial rise and diversification of angiosperms(38), suggesting that they are ecologically associated with fungi acting as saprotrophs or mycorrhizal partners (3). Finally, the age of mycenoid and marasmioid which was also the age of the luciferase cluster in fungi was estimated to originate around 160 million years ago during the late Jurassic (**Fig. 1B**).

### Differential conservation of synteny regions across *Mycena* genomes

We attempted to characterize chromosome evolution in the mycenoid clade using the newly available, highly contiguated assemblies for *Mycena*. We first compared the patterns of 4,452 single-copy orthologue pairs between assemblies of *Mycena indigotica* and *Armillaria ectypa* (*SI Appendix*, Fig. S8). The majority of scaffolds between the two species could be assigned one-to-one relationships unambiguously, providing strong evidence that macro-synteny has been conserved between the marasmioid and mycenoid clades. Such chromosome-level synteny remained conserved until the last common ancestor of the Agaricales, when *M. indigotica* was compared against the genome of *Pleurotus ostreatus* (*SI Appendix*, Fig. S9). Based on the clustering of single-copy orthologues, we identified 10 linkage groups resembling the number of known karyotypes in Basidiomycota suggesting possible ancestral chromosome numbers (25).

The *M. indigotica* scaffolds exhibit high orthologous gene density in the centres of scaffolds **(Fig. 2A)**. Fungal chromosomes can typically be compartmentalised into chromosomal cores and subtelomeres which display differential evolutionary dynamics (24, 39). In some extreme cases, filamentous pathogenic fungi contain entire lineage-specific chromosomes that are gene-sparse and enriched in transposable elements(40). In the case of *Mycena*, a multi-genome comparison showed that synteny conservation was typically either lost at the scaffold ends or extended by several mega-bases across the *Mycena* assemblies (**Fig. 3A**).

**Fig. 3.**
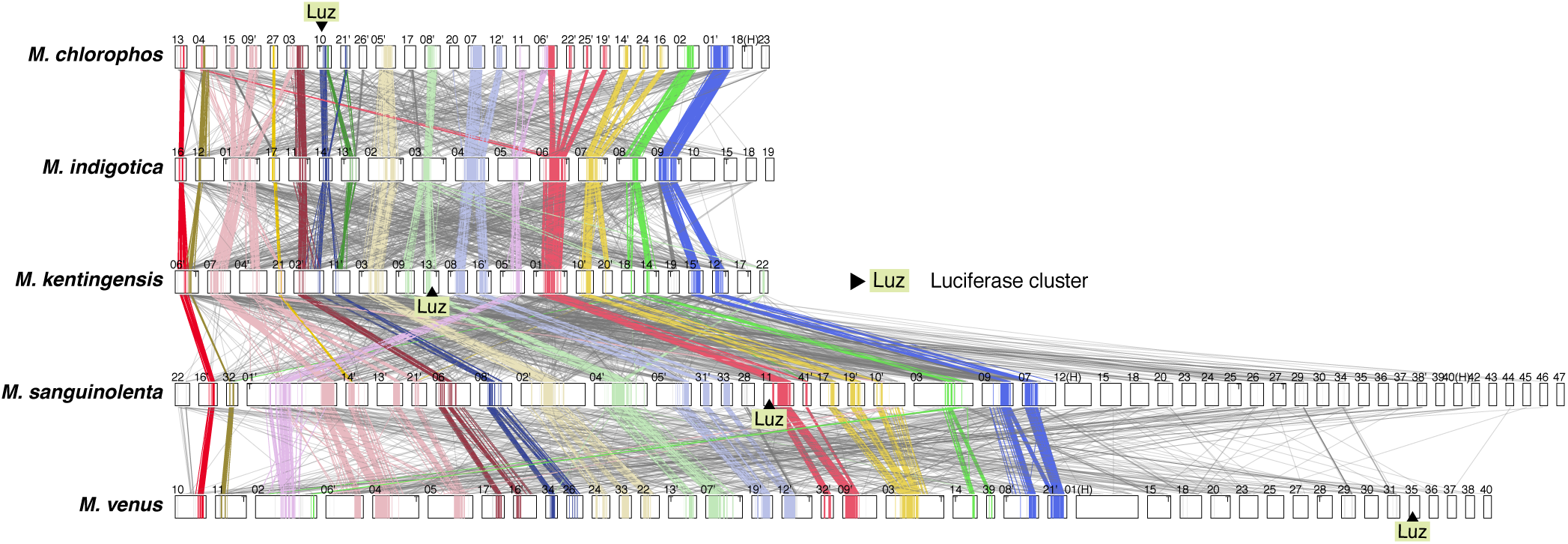
Genome synteny in *Mycena* genomes. Schematic representation of the inter-scaffold relationship between species. The lines between scaffolds denote single-copy orthologues between a pair of species. Shaded areas in each scaffold denote high-synteny regions defined by DAGchainer (41) and colour denote linkage groups assigned by most abundant pairwise single-copy orthologues. Lines are colour-coded according to corresponding linkage groups. Black triangles denote locations of luciferase clusters.

Defining precise boundaries between regions with and without synteny is challenging. Based on the clustering of orthologous genes using DAGchainer (41), we partitioned the scaffolds into low and high synteny regions. As expected, highly syntenic regions in *Mycena* were typically found at the scaffold centres. In contrast, synteny was not in parts of scaffold or, in some cases, throughout the entire scaffolds, as was the case for the largest (12.0 Mb) assembled scaffold of *M. venus* (**Fig. 3A**). These regions are highly enriched in repeats; they have 1.5–2.6-fold higher methylation levels and are overrepresented in expanded and contracted OGs compared to high synteny regions (*SI Appendix*, Fig. S10 and Table S10; two-proportions z-test, *P* < 2.1E-9). Expansions and contractions of gene families were 1.8–4.2 and 1.4–2.9 fold higher in the low than high synteny regions, respectively; differential gain and loss of genes in these regions may have important implications for *Mycena*.

### Evolutionary dynamics of luciferase clusters

One of the outstanding questions surrounding the evolution of fungal bioluminescence is why bioluminescent species are scattered across the mycenoid and marasmioid clades. The mechanism of fungal bioluminescence is homologous across species (6), and this implies that non-bioluminescent mycenoid and marasmioid species must have lost the functional luciferase gene cluster. To investigate the evolutionary dynamics of the luciferase cluster, we examined all highly contiguous assemblies across the bioluminescent lineages available and inspected adjacent synteny (**Fig. 4**). The majority of the *Mycena* luciferase clusters included luciferase (*luz*), hispidin-3-hydroxylase (*h3h*), cytochrome P450 (*cyp450*), and hispidin synthase (*hisps*). We found that physical linkage was only maintained within the luciferase cluster, and synteny was lacking in genes surrounding the luciferase cluster (**Fig. 4**) of species in the *Mycena* and *Omphalotus* lineages. Coupled with the aforementioned synteny analysis, we hypothesised that the luciferase cluster residing in a fast-evolving genomic region may result in it frequently being lost. The nearest TE sequence adjacent to luciferase cluster in *Mycena* species were 2-8.9kb away and separated by 0-5 genes suggesting possible roles of transposons mediating rearrangements (**Fig. 4**). Additionally, the luciferase cluster of different *Mycena* species was identified in low synteny regions and located in different linkage groups (**Fig. 3**), providing evidence that the location of the cluster had been extensively rearranged. In contrast, the genes surrounding the luciferase cluster among the eight *Armillaria* species were generally in the same order, with collinearity partially lost only in *G. necrorhiza* (a very close relative of *Armillaria*, **Fig. 4**). We found that the synteny surrounding the *Armillaria* luciferase cluster was maintained since the common ancestor of Agaricales (*SI Appendix*, Fig. S11). The up- and downstream regions of the luciferase cluster belonged to two separate regions of the same ancestral chromosome linkage group, suggesting that these regions were previously rearranged—including the luciferase cluster—and were subsequently retained (*SI Appendix*, Fig. S11).

**Fig. 4.**
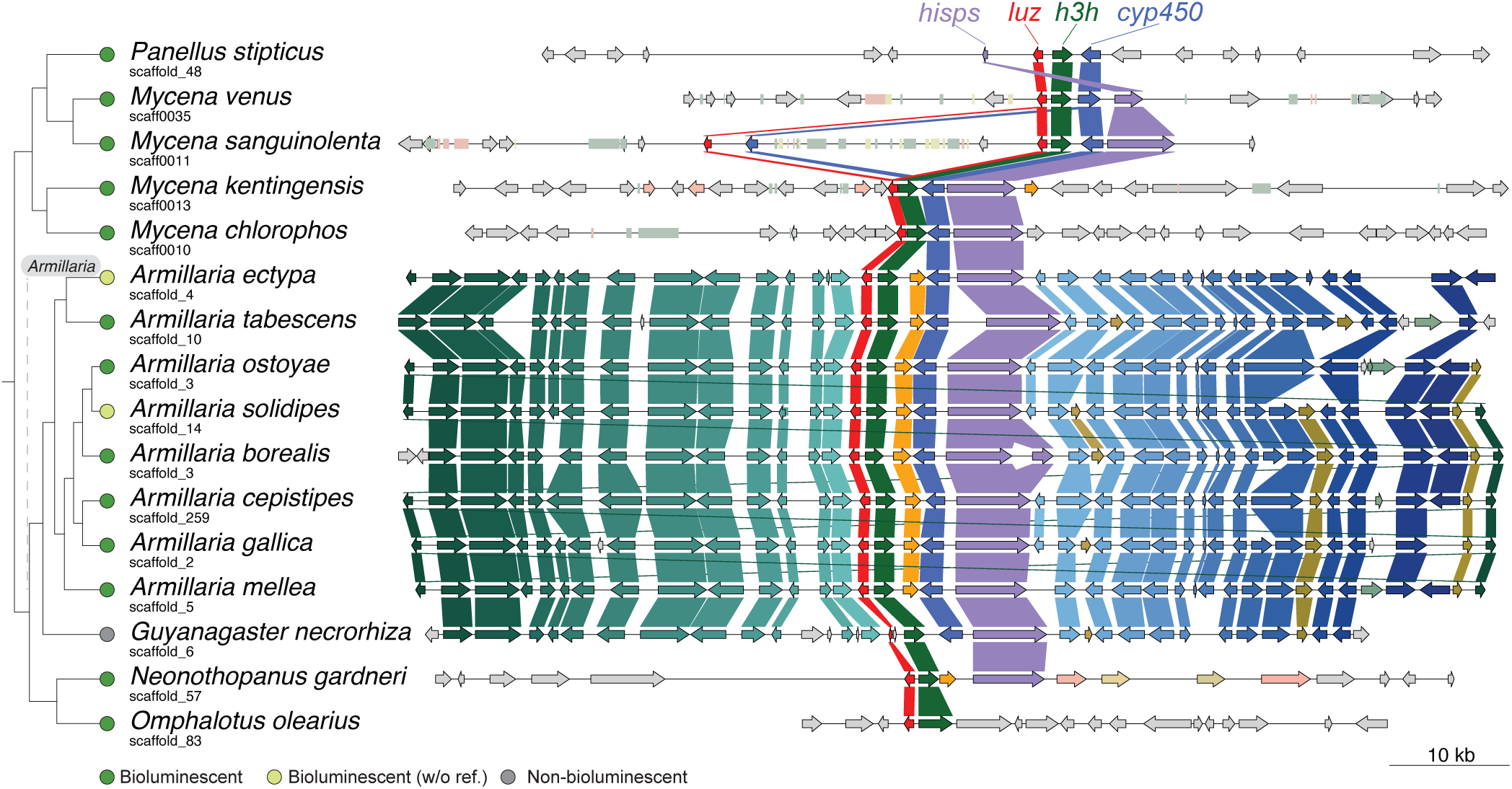
Synteny around the luciferase cluster among bioluminescent fungi. The orthologous groups (OGs) shared by at least two species were labelled with the same colour, regardless of their orientation. Arrows and rectangles denote protein encoding genes and transposable element, respectively. Different colours of rectangle denote TE types (pink: LINE and LINE relic; light green: LTR and LTR relic; yellow: DNA and DNA relic). The *cph* gene in some species was located in other scaffolds (Fig. S12).

These observations lead us to propose a most plausible evolutionary scenario in which the luciferase cluster evolved across all available bioluminescent fungi (**Fig. 5)**. We inferred that the ancestral luciferase cluster consisted of *luz, h3h, cyp450* and *hisps*, with caffeylpyruvate hydrolase *cph*—involved in oxyluciferin recycling(6, 7)—also present on the same chromosome. This combination was found in 14 of the 15 bioluminescent species used in this study. Our data outline two contrasting scenarios by which the luciferase cluster was retained. First, the luciferase clusters in family Physalacriaceae are located in slow-evolving chromosomal regions, resulting in all members of the *Armillaria* synteny retaining both many of the genes adjacent to the luciferase cluster and the uniform luciferase cluster in their genomes. By residing in slow-evolving regions, the luciferase cluster in *Armillaria* might not be prone to losses by frequent chromosomal events (rearrangements, TE activity, etc.), explaining why it is conserved in the genus. On the other hand, the luciferase cluster in the *Mycena* clade is located in a highly dynamic genomic partition with low synteny (**Fig 3**), which could explain why mycenoid fungi had a higher tendency to lose the luciferase cluster compared to *Armillaria* species.

**Fig. 5.**
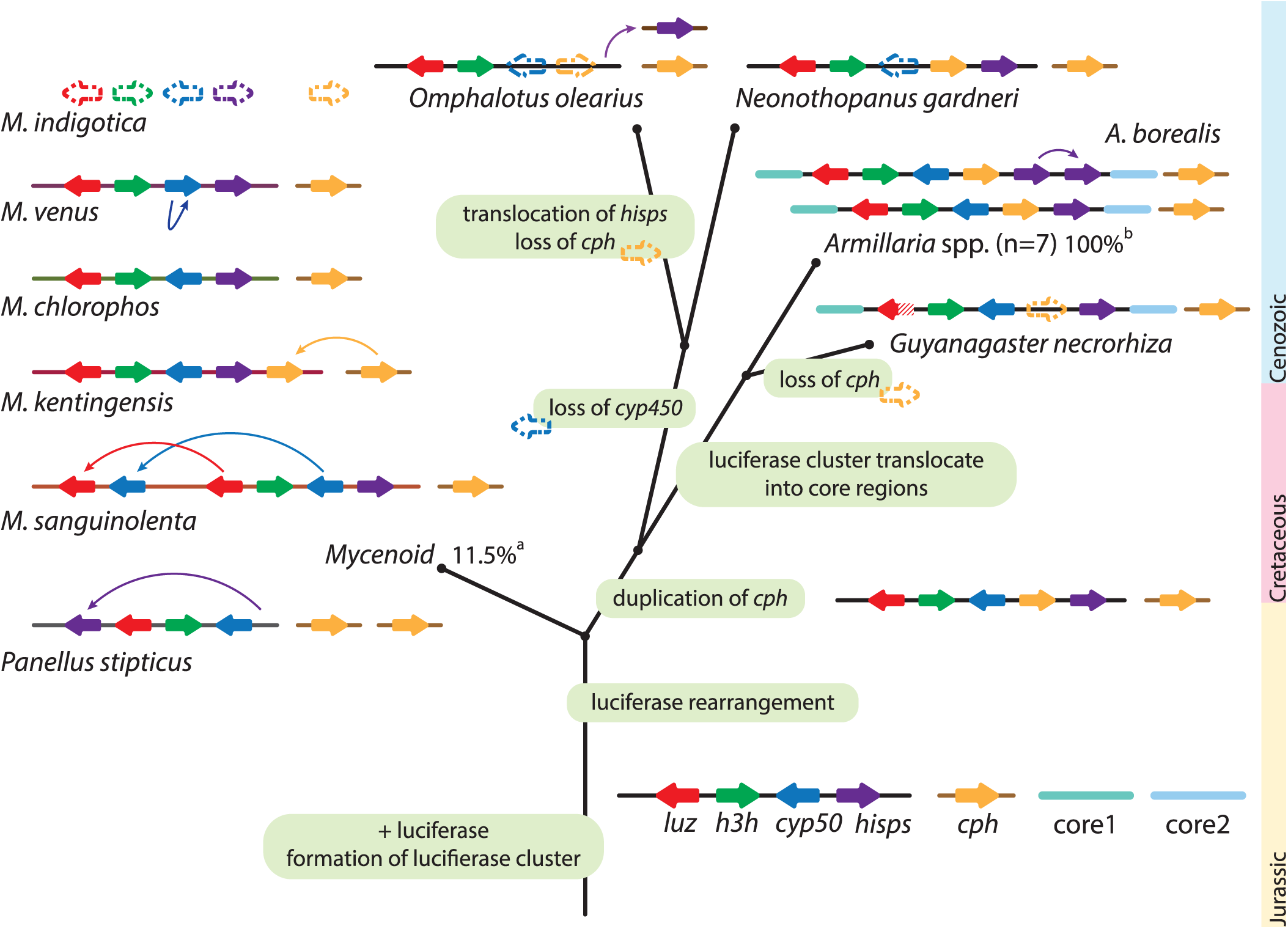
Evolutionary scenario for luciferase cluster evolution. The formation of the luciferase cluster originated at the dispensable region of the last common ancestor and was susceptible to translocate to different genomic locations through rearrangement. In the ancestor of marasmioid, *cph* was duplicated and translocated into the luciferase cluster. Before the ancestor of the Physalacriaceae family emerged, the luciferase cluster was translocated into the core region and have since kept its synteny in the *Armillaria* lineage. In the most recent common ancestor of *Mycena* species, the luciferase cluster was located in the dispensable region and have since been susceptible to further rearrangement. Arrow box indicates gene. The dashed arrow box denotes the loss of gene. Fishhook arrow denotes translocation event. ^a^ Percentage of bioluminescent fungi found in the mycenoid lineage(5). ^b^ Percentage of bioluminescent fungi found in *Armillaria* lineage(47).

Variations were common in the luciferase cluster. *cph* was located in different scaffolds in four of the five *Mycena* species (*SI Appendix*, Fig. S12). In *M. sanguinolenta, luz* and *cyp450* were duplicated adjacent to the luciferase cluster (**Fig. 4**). Losses were observed at different positions in the phylogeny. The non-bioluminescent *M. indigotica* lost the entire luciferase cluster, but *h3h* homologues were found in other regions of the genome, while *Guyanagaster necrorhiza* has a partial luciferase (7) and three other enzymes (**Fig. 4**), suggesting that an independent loss of luciferase function alone was enough for it to lose its bioluminescence. Interestingly, we found that the *cph* gene was independently translocated adjacent to the luciferase cluster in both *M. kentingensis* and the ancestor of the marasmioid clade (*SI Appendix*, Fig. S13); it was presumably favored and maintained here by natural selection (42). A selection analysis of genes in the luciferase cluster revealed that the majority of conserved sites exhibit either no or strong purifying selection, with only 7–28 sites under episodic selection (*SI Appendix*, Fig. S14). These results indicated that bioluminescence has limited roles in the species that have retained the process.

### Expression profile of luciferase cluster and identification of conserved genes involved in fungal bioluminescence

Fungal bioluminescence is believed to have ecological roles, such as attracting insects, and is regulated by circadian rhythms(43); however, the complete repertoire of genes involved in bioluminescence is still unknown. We carried out transcriptome profiling between mycelia with different bioluminescent intensities in four *Mycena* species, and identified genes that were either differentially expressed or positively correlated with bioluminescent intensities (Methods). There were 29 OGs found to contain upregulated gene members in all four *Mycena* species (**Fig. 1C and 6A**), including *luz, h3h*, and *hisps*, consistent with bioluminescence intensity dependent on the expression of these three genes in the luciferase cluster. In particular, *luz* expression was significantly different between two tissues with relative high and low bioluminescence in *M. kentingensis* (log fold change (logFC) 3.0; adjusted *P*< 0.001) and *M. chlorophos* (logFC 4.7; adjusted *P*<0.001); there was also a significant correlation between bioluminescent intensity and expression level in *M. sanguinolenta* (Pearson’s correlation coefficient (PCC) 0.82; *P*<0.005) and *M. venus* (PCC 0.86, *P*<0.005; Dataset S3. In *M. chlorophos*, however, its *cyp450* and *h3h* were not differentially expressed, and four distant homologues of *h3h* were found to be upregulated (*SI Appendix*, Fig. S7A). Although a second copy of *luz* and *cyp450* were found in *M. sanguinolenta*, they showed much lower expression (2 and 3 transcripts per million (TPM), respectively) than those in the cluster (282 and 138 TPM, respectively). The remaining OGs upregulated in mycelia showing higher bioluminescence included ABC transporters and Acetyl-CoA synthetases which also showed a predicted function in metabolic adaptations to bioluminescence in firefly and glowworm(44, 45). (**Fig. 6A**; *SI Appendix*, Table S11). In particular, four OGs were annotated as FAD or NAD(P)-binding domain-containing proteins. As these genes do not bear sequence similarity to *h3h* which is also a NAD(P)H-dependent enzyme, they are likely involved in other biochemical processes that is required during bioluminescence.

**Fig. 6.**
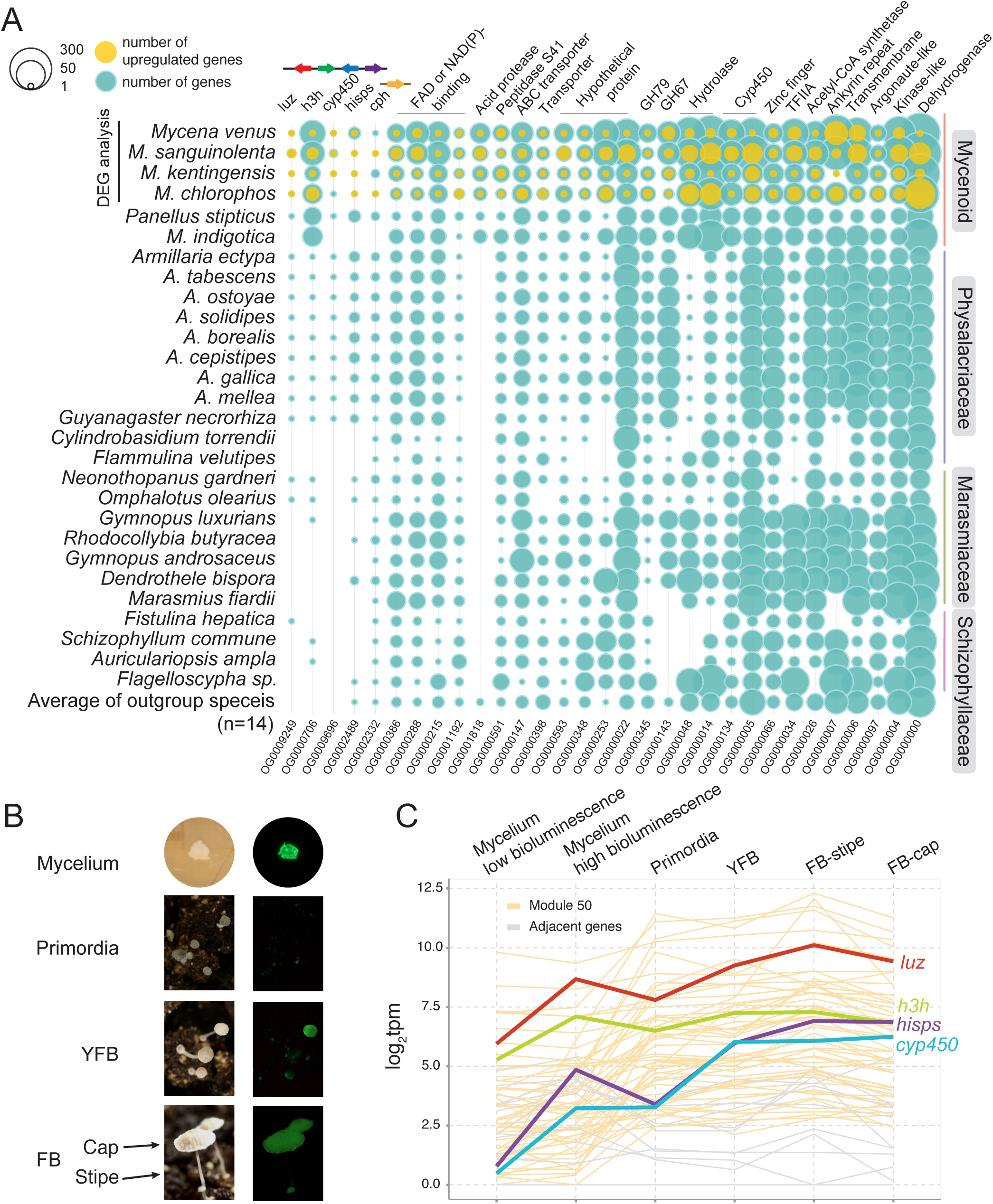
Expression analysis to identify genes involved in bioluminescence. (*A*) Conserved upregulated OGs. Differentially-expressed genes (DEGs) between mycelia with different bioluminescent intensities were identified in four bioluminescent *Mycena* species, and all 29 OGs—except OG0009249 and OG0000706—contain at least one upregulated gene. A detailed annotation of the genes in the OGs is listed in Table S11. (*B*) Tissues used for transcriptomic data analysis in *M. kentingensis*. The left and right side are the tissues under light and dark conditions, respectively (captured by a Nikon D7000). The camera setting for each tissue: mycelium, Sigma 17-50mm ISO100 f2.8 with 16 min exposure time; primordia, AF-S Micro Nikkor 60mm ISO800 f/11with 122.4 sec exposure time; YFB, AF-S Micro Nikkor 60mm ISO800 f/11with 60.6 sec exposure time; FB, AF-S Micro Nikkor 60mm ISO800, f/11 with 9.3 sec exposure time. YFB, young fruiting body (0.5-1 cm). FB, mature fruiting body (> 1 cm). FB-cap, cap from FB. FB-stipe, stipe from FB. (*C*) Expression profile of luciferase cluster across developmental stages of *M. kentingensis*. Bold lines indicate four genes in the luciferase cluster. These four genes and the other 53 genes (yellow) were assigned into the same module (Module50) with similar expression patterns. The genes located up- or downstream (grey) of the luciferin biosynthesis cluster had lower expression levels than the four genes in the cluster.

Differences in bioluminescent intensity have been recorded in tissues of fungi both in nature (4, 5, 46-48) and—for *M. kentingensis—*in a laboratory environment, in which the life cycle can be completed (**Fig. 6B**). To investigate putative roles of bioluminescence across developmental stages, additional transcriptome profiling was carried out in the primordia, young fruiting body, and cap (pileus) and stipe of the mature fruiting body of *M. kentingensis*. Bioluminescence was stronger in the cap than in the stipe, so we expected the luciferase cluster genes to have higher expression in the cap tissue. However, *luz* and *h3h* showed opposite expression patterns (**Fig. 6C** and Dataset S4), suggesting that there may be other regulators involved in bioluminescence in *M. kentingensis*.

The regulation of bioluminescence in *M. kentingensis* during development was determined by performing a weighted correlation network analysis (WGCNA(49, 50)), which identified 67 modules of co-expressed genes in these stages (*SI Appendix*, Fig. S15). All members of the luciferase cluster *luz, h3h, cyp450*, and *hisps* belonged to the same module (Module50; **Fig. 6C**) of 57 genes, suggesting that the expression of the luciferase cluster members are co-regulated during developmental stages. Only two genes belonging to OG0001818 (acid protease) and OG0000000 (short-chain dehydrogenase) which were part of the 29 aforementioned OGs associated with bioluminescence in the mycelium samples across *Mycena*. Six genes in this module were annotated as carbohydrate-active enzymes (Dataset S5): one GH75 (chitosanase), one AA1_2 (Laccase; ferroxidases), two GH16, and two genes with two CBM52 domains. GH16 (glucanases) and AA1 (laccases) are known to be differentially expressed during fruiting body development(51), implying a possible link between cell wall remodelling during development and bioluminescence. In addition, we re-analysed the transcriptomes of *Armillaria ostoyae* across different developmental stages from Sipos *et al*. (2017)(9). Consistent with the observation that bioluminescence was only observed in mycelia and rhizomorphs in *A. ostoyae*(52, 53), the expressions of *luz, h3h, cyp450*, and *cph* were highest in these tissues (*SI Appendix*, Fig. S16). Together, these results imply that the luciferase cluster was differentially regulated during development and that the extent of the expressions was also different among bioluminescent species of different lineages.

### Gene families associated with the evolution of mycenoid species

We assessed orthologous group evolution by analysing OG distribution dynamics along a time-calibrated phylogeny using CAFÉ (54). The rate gene family changes in mycenoid were comparable to those of other branches of Agaricales (likelihood ratio test; *P* = 0.25). A total of 703 orthologous groups were expanded at the origin of the mycenoid lineage (*SI Appendix*, Fig. S17). Analysis of gene ontology terms showed that these genes were enriched in NADH dehydrogenase activity, monooxygenase activity, iron ion binding, and transferase activity (Dataset S6). Additionally, we sought to identify proteins specific to mycenoid species by annotating protein family (Pfam) domains and comparing them with those of species outside this lineage (Dataset S7). A total of 537 Pfam domains were enriched in the mycenoid lineage (one-fold by Wilcoxon rank sum test with *P*<0.01; Dataset S8) of which 3–17 were species-specific. Acyl_transf_3 (acyltransferase family; PF01757), contained in a range of acyltransferase enzymes, was the only domain found in all six mycenoid species. The closest homologs were found in ascomycetous *Cadophora, Pseudogymnoascus*, or *Phialocephala* (31-35% identity with 73-100% coverage). Four of the enriched domains are known pathogenesis-related domains expanded in pathogenic Agaricales *Moniliophthora* (55) and *Armillaria* species(9): COesterase (PF00135; Carboxylesterase family), Thaumatin (PF00314), NPP1 (PF05630; necrosis-inducing protein), and RTA1 (PF04479; RTA1-like protein) (*SI Appendix*, Fig. S18). Moreover, *M. sanguinolenta* and *M. venus* contained over 100 and 17 copies of COesterase and Thaumatin (median 37 and 4 copies in other fungal species of this study), respectively.

## Discussion

Bioluminescence is one of the most unusual and fascinating traits in fungi, but the evolutionary history of the luciferase gene cluster, which underlies this phenomenon, has remained elusive. Here, we produced highly contiguous genome assemblies using Nanopore technology and annotations for five of the *Mycena* species to examine their genome dynamics and bioluminescence. The results of phylogenomic analyses on these genomes have important implications for the origin of luciferases.

The first question we addressed is whether fungal bioluminescence originated once or multiple times. Our species phylogeny is in good general agreement with comparative genomic analyses around this group(9). We show that the fungal luciferase, which represents a *de novo* origin of luciferase activity different from that in other lineages (insects, bacteria, etc), first appeared together with other members of the luciferase cluster in the last common ancestor of mycenoid and marasmioid clades. Compared to previous inferences that this cluster had a single origin, our results imply extensive loss of the luciferase cluster in these two clades (**Fig. 1B** and **1C**) to explain the patchy phylogenetic distribution of and minor presence for bioluminescence in fungi. An alternative scenario could be that fungal bioluminescence arose multiple times through convergent evolution. Although multiple origins yield a more parsimonious model, we can confidently reject this hypothesis because both the fungal luciferase and the luciferase gene cluster are clearly homologous across distant bioluminescent fungi (6). Although a single origin and the excessive number of implied losses may appear counterintuitive, models of trait evolution and recent empirical evidences of phylogenetically patchy, but homologous traits (56, 57) have emerged offering a biologically reasonable explanation. One attractive model is the latent homology model (58), which posits that precursor traits can potentiate lineages for easily and recurrently evolving similar traits. More comprehensive surveys of genomes in these lineages are needed to make informed speculations on whether latent homologies might have facilitated the evolution of bioluminescence in fungi.

The next outstanding question is therefore what caused the frequent losses of bioluminescence in fungi? Our evolutionary reconstructions show that the luciferase cluster might have originated in low-synteny region of genomes (**Fig. 3**), making it susceptible to rearrangement, which suggests it is highly prone to loss and explains why most mycenoid and marasmioid species are non-bioluminescent. This is consistent with a previous report that the main evolutionary process in fungal gene clusters is vertical evolution followed by differential loss (59). Interestingly, synteny was retained in luciferase clusters and adjacent genes of *Armillaria* species (**Fig. 4**), which are better known for their roles as plant pathogens(9). Such synteny remained detectable when compared to representative Agaricales and *P. ostreatus* genomes, suggesting that in *Armillaria*, the luciferase cluster was translocated to a region of the genome where synteny was conserved. Indeed, bioluminescence was identified in all nine examined *Armillaria* species(47). The alternative but less parsimonious scenario would be that the luciferase cluster originated in a high-synteny region and subsequently translocated to low-synteny regions in ancestors of both mycenoid and various families within the marasmioid clade. The repeated duplication and relocation of *cph* that we observed in the luciferase cluster is under selection pressure, suggesting that bioluminescence was maintained in fungi that still exhibit this phenotype. A systematic quantification of bioluminescence and more complete genome assemblies will help reconstruct the evolutionary events that contributed to the polymorphism and functional diversity in the luciferase clusters.

Researchers have long been puzzled over the ecological role of bioluminescence in fungi. One explanation that has been put forth for *Neonothopanus gardneri* is that bioluminescence follows a circadian rhythm to increase spore dispersal by attracting arthropods in the evening(43). If true, this is most likely a derived adaptation, as most bioluminescent fungi — including *Mycena, Omphalotus* and *Armillaria* species — disperse spores via wind, display bioluminescence continuously, and are not known to attract insects(60). Besides, attraction is insufficient to explain luminescence in the mycelium. We have shown that the luciferase cluster in *Mycena kentingensis* is constitutively expressed throughout development. We further identified a handful of genes whose expressions are correlated with fungal bioluminescence and may therefore be candidates for experimental follow-up studies (**Fig 6**). If fungal bioluminescence originated as a by-product of a biological process that is currently unknown, the ecological role was likely to be initially limited which may explain why it has undergone subsequent losses in many species. For those that have retained bioluminescence, its ecological role remains unknown, but we speculate that it may be species-specific, explaining why the luciferase cluster had been maintained across hundreds of millions of years.

In summary, our comparative analyses allowed us to propose an evolutionary model pinpointing changes in the luciferase cluster across all bioluminescent fungi with published genomes. Our findings offer insights into the evolution of a gene cluster spanning over 160 million years and suggests that the retained luciferases were under strong purifying selection. Our *Mycena* genome sequences may complement ongoing research on the application of bioluminescent pathways(7) and shed light on the ecological role of bioluminescence in fungi.

## Methods

More detailed information on the materials and methods used in this study are provided in *SI Appendix*.

### *De novo* assemblies of *Mycena* species

Haploid genome length and heterozygosity of the five *Mycena* species (*M. kentingensis, M. venus* (61), *M. sanguinolenta, M. indigotica* and *M. chlorophos)* were estimated from Illumina reads using GenomeScope (14) (ver. 2.0). Oxford Nanopore reads were assembled using the Canu(13) (ver. 1.8) assembler. Consensus sequences of the assemblies were polished first by five iterations of Racon(62) (ver. 1.3.2) followed by Medaka (ver. 0.7.1; https://github.com/nanoporetech/medaka) using Oxford Nanopore reads. HaploMerger2(63) (ver. 20180603) was then run on to generate haploid assemblies. Finally, the consensus sequences were further corrected with Illumina reads using Pilon(64) (ver. 1.22). Quality values (QV) of the final assemblies were calculated as described in Koren *et al* (65). Throughout each stage the genome completeness was assessed using fungi and basidiomycete dataset of BUSCO(19) (ver. 4.1.2). Putative telomeric repeats were searched for copy number repeats less than 10 mers using tandem repeat finder(66) (ver. 4.09; options: 2 7 7 80 10 50 500). The hexamer TTAGGG was identified (*SI Appendix*, Table S12).

### Gene predictions and functional annotation

Protein sequences from Uniprot fungi (32,991 sequences; downloaded 20^th^ December 2018) and *Coprinopsis cinerea, Pleurotus ostreatus* PC15 (v2.0), *Schizophyllum commune and Armillaria mellea* from MycoCosm(67) portal were downloaded as reference proteomes. Transcriptome reads were first mapped to the corresponding genome assemblies using STAR(68, 69) (ver. 2.5.3a), and subsequently assembled into transcripts using Trinity(70) (ver. 2.3.2; guided approach), Stringtie(71) (ver. 1.3.1c), CLASS2(72) (ver. 2.1.7) and Cufflinks(73) (ver. 2.2.1). The samples used for input are listed in Dataset S1. Transcripts generated from Trinity were aligned to the references using GMAP(74). All transcripts were merged, filtered and picked using MIKADO(75) (ver. 1.1). The gene predictor Augustus(76) (ver. 3.2.1) and gmhmm(77) (ver. 3.56) were trained using BRAKER2(78) (option fungi and softmasked), and SNAP(79) was trained using the assembled transcripts with MAKER2(18) (ver. 2.31.9). The assembled transcripts, reference proteomes and BRAKER2 annotations were combined as evidence hints for input in the MAKER2(18) annotation pipeline. MAKER2(18) invoked the three trained gene predictors to generate a final set of gene annotation. Descriptions of amino acid sequences of the proteome were annotated using Blast2GO(80) and GO terms were annotated using Argot (ver.2.5; (81). 80.0–86.7% of proteomes were assigned at least one GO term. Genes encoding carbohydrate-active enzymes were identified using dbCAN (82) (database ver. Hmm9.0; code ver. 2.0.0) by searching for sequence homologs with HMMER (83). Consensus (library) sequences of repetitive elements were identified using the pipeline described in Berriman *et al*(84).

### Methylation analyses

High-quality paired-end reads were aligned to the genome assemblies of *M. kentingensis* using the bisulfite specific aligner BS-Seeker2(85). Only uniquely mapped reads were retained. The cytosines covered by at least four reads were included in the data analysis, and the DNA methylation level for each cytosine was estimated as #C/(#C+#T), where #C is the number of methylated reads and #T is the number of unmethylated reads.

One or two Nanopore flowcells for each *Mycena* species were selected to infer methylation information using deepsignal (26) (ver. 0.1.5) (*M. kentingensis*: FAH31207, *M. chlorophos*: FAH31470, *M. indigotica*: FAH31228, *M. sanguinolenta*: FAK22405 and FAH31211, *M. venus*: FAK22389 and FAH31302). The machine learning-based model was trained with one bisulfite dataset (YJMC0389) and one Nanopore dataset (FAH31207) of *M. kentingensis*. The bisulfite result was first filtered for depth >20, then methylation levels >0.9 and <0.01 were selected for positive and negative validation datasets, respectively. All seven flowcells were called for methylation information with a customized model and default arguments. A minimal depth of 4 was applied to the results for further analysis. In the estimates of DNA methylation levels between Nanopore long-reads and the Illumina BS-seqs, the Pearson correlation coefficient was as high as 0.96 in the methylomes of *M. kentingensis* (*SI Appendix*, Fig. S19).

### Phylogenomic analyses

Orthologous groups (OGs) among 42 species were identified using OrthoFinder(20, 21) (ver. 2.2.7). A total of 42 sets of amino acid sequences from 360 single-copy OGs were aligned independently using MAFFT(86) (ver. 7.271; option --maxiterate 1000). A total of three approaches were used to infer the species tree. The first two approaches relied on maximum likelihood phylogenies from individual OG alignments computed using RAxML-ng(87) (ver. 0.9.0; options: --all --model LG+I+F+G4 --seed 1234 --tree pars 10 --bs-trees 100) with 100 bootstrap replicates. The best phylogeny and bootstrap replicates were separately used to infer a consensus tree using ASTRAL-III(36). Finally, a maximum likelihood phylogeny from the concatenated amino acid alignments of the single-copy orthogroups was constructed with 100 bootstrap replicates using RAxML-ng(87) (ver. 0.9.0; options: --all --seed 1234 --tree pars 10 --bs-trees 100 with --model LG+I+F+G4 partitioned with each OG alignment).

### Estimation of divergence time

The divergence time of each tree node was inferred using MCMCtree in PAML(37) package (ver. 4.9g with approximate likelihood(88); the JC69 model and the rest were default). The individual amino acid alignments of 360 single-copy-orthologs were converted into corresponding codon alignments using PAL2NAL (89) (ver. 14). The species tree and concatenated alignments of these single-copy-orthologs were used as the input for MCMCtree. The phylogeny was calibrated using fossil records by placing soft minimum bounds at the ancestral node of: i) marasmioid (using *Archaeomarasmius*□*legettii* 94–90 Ma(90); 90 was used), ii) Agaricales (using *Palaeoagaricites*□*antiquus* 110–100 Ma(91); 100 was used), iii) Taxon A (∼99 Ma(92); 95 was used), and iv) a soft bound of 200 Ma for the phylogeny. The entire analysis was run five times to check for convergence.

### Synteny analyses

Linkage groups (LGs) between *M. indigotica* and *Armillaria ectypa*, and between *M. indigotica* and *Pleurotus ostreatus* were assigned based on the reciprocal majority of the single-copy orthologues (*SI Appendix*, Fig. S8 and S9). Scaffolds that contained fewer than 10 single-copy orthologues, shorter than 500 kb or shorter than species N90 were excluded from the analysis. Linkage groups within *Mycena* were assigned based on majority and at least 10% of single-copy orthologue links with *M. indigotica* scaffolds. Subsequent scaffolds were identified as the same linkage group if they contained a majority of pairwise one-to-one single-copy orthologues belonging to the *M. indigotica* LG.

As gene collinearity among *Mycena* species became less conserved, synteny blocks of each *Mycena* species were defined based on merging of adjacent pairwise single-copy orthologues to its closest-related species. For instance, synteny blocks of *M. chlorophos* were based on single-copy orthologues against *M. indigotica*. For every ortholog, the distance to the next closest single-copy orthologue was calculated to take into account segment duplications of genes or gene insertion/deletions. Synteny blocks of each species were estimated from pairwise proteome comparisons against its closest relative using DAGchainer (41) (options -Z 12 -D 10 -g 1 -A 5). Synteny around luciferase cluster was plotted using the genoPlotR(93) package.

### RNAseq analysis of differential bioluminescent mycelium

Quality trimming of the RNA sequencing reads was conducted using Trimmomatic(94). The sequencing reads were mapped to the genome using STAR(68, 69) (ver. STAR_2.5.1b_modified; default parameters). Raw read counts of the gene models were quantified by FeatureCounts(95) (ver. v1.5.0; -p -s 2 -t exon). For *M. kentingensis* and *M. chlorophos*, the differential expressed genes (DEGs) were analysed using DESeq2(96). Genes with fold change (FC) > 0 and FDR ≤ 0.05 were defined as DEG. *For M. sanguinolenta* and *M. venus*, the DEGs were identified by the Pearson correlation coefficient between the bioluminescence intensity (relative light unit; RLU) normalized by weight (RLU/mg) and log transformation of counts per million. Genes with correlation coefficient > 0.7 and *P*<0.01 were defined as DEGs.

### RNA analysis of *M. kentingensis* and *Armillaria ostoyae* developmental stages

The reads from transcriptomes of the primordia, young fruiting body, and cap and stipe of mature fruiting body were conducted by the same method of manipulating the reads from transcriptomes of mycelium. To identify co-expressed genes among transcriptomes, the transformation of transcripts per million (TPM) from six different tissues—mycelia with high bioluminescence and low bioluminescence, primordia, young fruiting body, and fruiting body cap and stipe were calculated. The lowest 25% expressed gene across all samples were excluded and co expression was analysed using weighted gene co-expression network analysis (WGCNA)(49, 50) package in R (maxBlockSize = 10000, power = 20, networkType = signed, TOMType = signed, minModuleSize = 30). The Illumina reads among ten stages from *Armillaria ostoyae* were also downloaded from NCBI’s GEO Archive (http://www.ncbi.nlm.nih.gov/geo <u>under accession GSE100213</u>) and also analysed by the same pipeline of *M. kentingensis* to identify co-expressed genes among the transcriptomes.

## Authors contribution

I.J.T. and H.M.K. conceived the study. I.J.T. led the study. H.M.K., C.C.C., G.S. and H.W.K. collected and identified *Mycena* species around Taiwan. H.M.K, P.H.W. and C.I.L. conducted the experiments. M.J.L. and J.Y.L. designed the Illumina sequencing experiment. H.H.L. and I.J.T. performed the assemblies and annotations of the *Mycena* genomes. H.M.K., H.H.L., Y.C.L. and I.J.T. conducted the repeat analysis. L.G.N. and I.J.T. carried out phylogenomics analyses and the divergence time estimation. H.M.K., H.H.L., Y.C.L. and I.J.T. carried out comparative genomic analyses. H.M.K. and M.R.L. analysed the RNA-seq data. H.H.L., R.J.L., J.W.H., P.Y.C. and H.M.K. carried out the methylation analyses. H.M.K. and I.J.T. wrote the manuscript with input from L.G.N and P.Y.C.

## Acknowledgement

We thank Chia-Ning Shen for lending us the luminometer for the duration of the project. We thank Bi-Chang Chen for providing the CCD camera to qualify bioluminescence. We thank Chi-Yu Chen and Jie-Hao Ou for their useful advice on culturing *Mycena* fungi. We are grateful to the National Center for High-performance Computing for computer time and facilities. We are grateful to the ‘1000 Fungal Genomes – Deep Sequencing of Ecologically-relevant Dikarya’ consortium for access to unpublished genome data. We thank Gregory Bonito, Hui-Ling Liao, Alejandro Rojas and Rytas Vilgalys for permission to use the *Flagelloscypha* sp. FlaPMI526_1 assembly from JGI. We thank Mary Catherine Aime for permission to use the *G. necrorhiza* MCA 3950. assembly from JGI. The genome sequence data were produced by the US Department of Energy Joint Genome Institute (JGI) in collaboration with the user community. We thank Chia-Lin Chung, Ben-Yang Liao and John Wang for commenting the earlier version of the manuscript. P.Y.C was supported by Ministry of Science and Technology, Taiwan, under Grant No. 106-2311-B-001 −035 -MY3 and 108-2313-B-001 -013 -MY3. H.M.K was supported by postdoctoral fellowship, Academia Sinica. I.J.T was supported by Career Development Award AS-CDA-107-L01, Academia Sinica. L.G.N was supported by the ‘Momentum’ Program of the Hungarian Academy of Sciences (grant # LP2019-13/2019).

## Data availability

Genome assembly and annotation of five *Mycena* species was deposited in the National Centre for Biotechnology Information BioProject database (accession no. PRJNA623720) pending final checks.

