## Supplementary Information for "*Mycena* genomes resolve the evolution of fungal bioluminescence"

Isheng Jason Tsai

Huei-Mien Ke and Isheng Jason Tsai

**This PDF file includes:**

Supplementary text

Figures S1 to S19

Tables S1 to S13

Legends for Datasets S1 to S9

References for SI reference citations

**Other supplementary materials for this manuscript include the following:**

Datasets S1 to S9

**I. SUPPLEMENTARY INFORMATION TEXTS**

**Materials and Methods**

**Strains and fungal materials**

*M. kentingensis*, *M. venus* (1)*, M. sanguinolenta, M. indigotica* and *M. chlorophos* were isolated from fruiting bodies collected from forest in Taiwan. *M. indigotica* was isolated from basidiospores. The mycelia were grown and maintained on potato dextrose agar (PDA) plates at 25°C. To identify the pattern of bioluminescence, a piece of mycelium from each species was inoculated in the centre of a sheet of sterilized dialysis cellulose membrane (8030-32, Cellu Sep T-Series) on a 3 cm PDA agar plate at 25°C. The diameter of the mycelium was measured and its bioluminescence was recorded with a Glomax 20/20 luminometer (Promega BioSystems Sunnyvale, Inc., USA) for seven days (Dataset S9). The taxonomic status of species was reconfirmed by sequencing the internal transcribed spacer (ITS) with the primer pair SR6R(5’-AAGWAAAAGTCGTAACAAGG-3’)/ITS4(5’-TCCTCCGCTTATTGATATGC-3’). Using the other available *Mycena* ITS sequences, all sequences were aligned by MAFFT(2) (ver. 7.310) and trimmed by trimAl(3) (1.2rev59; with option -automated1). The ITS phylogeny was constructed by IQ-TREE(4, 5) (ver. 1.6.10; with option -bb 10000 -alrt 1000).

**Genomic DNA extraction and sequencing**

Genomic DNA was extracted using the traditional CTAB and chloroform extraction method. Briefly, 0.1 g mycelium was grinded with liquid nitrogen and then mixed with CTAB extraction buffer (0.1 M tris, 0.7 M NaCl, 10 mM EDTA, 1% CTAB, 1% Beta-Mercaptoethanol). After incubating at 65°C for 30 min, an equal volume of chloroform was added, then the mixture was centrifuged at 8000 rcf for 10 min. The supernatant was mixed with an equal volume of isopropanol and the DNA was precipitated. After washing with 70% EtOH, the DNA was dissolved with nuclease free water. Genome sequencing was carried out in two platforms. First, paired-end libraries were constructed using the KAPA LTP library preparation kits (#KK8232, KAPA Biosystems). All libraries were prepared in the High Throughput Genomics Core at Biodiversity Research Center, Academia Sinica and sequenced on an Illumina HiSeq 2500 platform. A total of 51.6 Gb of 150- or 300-bp read pairs were generated. Second, Oxford Nanopore libraries were prepared using SQK-LSK108 and sequenced on a GridION instrument. Basecalling of Nanopore raw signals was performed using Guppy (ver. 3.2.4) into a total 67.7 Gb of raw sequences at least 1 kb or longer. A summary of the sequencing data is shown in *SI Appendix,* Table S1.

**RNA extraction and sequencing**

Bioluminescent mycelia were collected in two ways. i) For *M. chlorophos* and *M. kentingensis*, a piece of mycelium was inoculated at the centre of a sheet of sterilized dialysis cellulose membrane (8030-32, Cellu Sep T-Series) on PDA agar plates at 25°C. The plates were cultured for 10 and 14–18 days for *M. chlorophos* and *M. kentingensis*, respectively. For *M. kentingensis*, bioluminescence was detected by camera (Nikon D7000, Sigma 17-50mm ISO100 f2.8 with 16 min exposure time) (**Fig. 6B**). The mycelia with low or high bioluminescent intensities which occurred spontaneously were collected from two separated plates inoculated on the same day. In *M. chlorophos*, bioluminescence was detected by luminometer. Mycelium with low bioluminescence showed the intensity of 7-14 Relative Light Unit (RLU)/mg, and the mycelium with high bioluminescence showed the intensity of 5,000-10,000 RLU/mg (Dataset S1). Three replicates were collected. ii) For *M. sanguinolenta* and *M. venus*, a piece of mycelium was inoculated at the centre of a sheet of sterilized dialysis cellulose membrane on PDA agar plates at 25°C. The plates were cultured for 13–17 days, and the bioluminescent features were detected by CCD camera; the tissues were collected and their luminescence intensity was recorded with a Glomax 20/20 luminometer (Promega BioSystems Sunnyvale, Inc., USA). A total of 12 samples with different bioluminescence intensities were collected (Dataset S1). After homogenizing 5–10 mg of tissues by liquid nitrogen, total RNA was extracted using the Direct-zol RNA Miniprep (Zymo Research). Concentrations were measured by Qubit fluorometer (Invitrogen USA), and quality was assessed by the BioAnalyzer 2100 RNA Nano kit (Agilent, USA) with RIN values higher than 8.0. The paired-end libraries were constructed using the TruSeq Stranded mRNA library prep kit (#20020594, Illumina, San Diego, USA) with standard protocol and sequenced by Illumina HiSeq 2500 (Illumina, USA) to produce 150-bp paired-end reads.

**RNA extraction and sequencing from the *M. kentingensis* fruiting body**

Fruiting body production of *M. kentingensis* was modified from previous studies(6, 7). Mycelia, grown on PDA for 8–15 days, was then inoculated onto sterilized commercially available peat soil mixed with 10% rice bran and 50% water in a jar. Mycelium samples were grown at 25°C for 3–4 weeks and then transferred into fresh compost. The culture was sprayed with sterilized water daily until the fruiting body formed. Four kinds of tissue were collected: (1) primordia, (2) young fruiting body (YFB, 0.5–1 cm), (3) cap and (4) stipe of mature fruiting body (> 1 cm). For each batch of culture, 15–20 primordia, 6–11 YFB, and 8–12 caps and stipes from mature fruiting bodies were pooled to measure their weight and bioluminescent intensity, and the RNA was extracted using Trizol extraction and lithium chloride purification method. Three replicates were produced. The paired-end libraries were constructed using the TruSeq Stranded mRNA library prep kit (#20020594, Illumina, San Diego, USA) with standard protocol and sequenced by Illumina HiSeq 2500 (Illumina, USA) to produce 150-bp paired-end reads.

**Bisulphite sequencing**

To construct a BS-seq library, the fragmented DNA was first ligated with a premethylated TruSeq DNA adapter (Illumina). The ligated DNA fragments were bisulfite converted using the EZ DNA methylation kit (Zymo Research), followed by PCR amplification. The BS-seq libraries were sequenced on an Illumina HiSeq 2500 sequencer. The bisulfite conversion efficiency reached approximately 99% in all of our libraries (*SI Appendix,* Table S13).

**Identification of repetitive elements**

Consensus (library) sequences of repetitive elements were identified using the pipeline described in Berriman *et al*(8). Full LTR retrotransposons in *Mycena* species were defined as i) initially identified by LTRharvest(9) and ii) presence of known reverse transcriptase domains identified by Pfam(10) (ver. 31.0). Repeat contents were quantified using RepeatMasker(11) (ver. open-4.0.7). Proportions of repeat content along the scaffolds were calculated using Bedtools(12). A phylogenetic tree was built by first aligning all the putative RVT domain sequences using MAFFT(2) (ver. 7.310; --genafpair --ep 0) and FastTree(13) with the JTT model on the aligned sequences, and were visualised using the ggtree(14) package in R.

**Orthogroup inference and analysis of protein family domains**

CAFÉ(15) (ver. 4.2.1; lambda command) was used to predict the expansion and contraction of gene numbers of OGs based on the topological gene tree. Gene family evolutionary rate λ was estimated for the whole phylogeny as well as a separate λ since the last common ancestor of the mycenoid lineage. Simulated dataset (option -t 120) were created by the genefamily command using the two different λ and significance was assessed using a likelihood-ratio test. The phylogenetic tree was visualized by the ggtree(14, 16) package in R. Protein domains of each gene were identified by pfam_scan.pl ver. 1.6 by comparing them against Pfam ver. 32.0 db(10). To compare them to plant pathogenic fungi, the Pfam domains from *Moniliophthora perniciosa* FA55313 (Monpe1_1)(17) from JGI and *Moniliophthora roreri* (Monro) from BioProject: PRJNA279170 were also annotated. Enrichment of Pfam domain number between two sets of interest was assessed by the Wilcoxon rank-sum test (*P* ≤ 0.05). We compared the Pfam copy number between six mycenoid species and the other 37 species. Gene ontology enrichments were identified for these genes using TopGO(18).

**Evolution of gene families related to the luciferase gene cluster**

In addition to gene family identified by Orthofinder (19, 20), luciferase orthologue outside marasmioid+mycenoid were identified by the reciprocal best hits against proteomes of three bioluminescence fungi (*Armillaria mellea*, *Neonothopanus gardneri*, and *Mycena kentingensis*). Outgroup sequences for the luciferase phylogeny was identified by first producing a phylogeny of these luciferase sequences with top 100 non-redundant hit of *M. sanguinolenta* (Msan_01367500) or *Fistulinahepatica* (Fishe1_70153) luciferase sequence. The sequences were aligned by MAFFT (2) (ver. 7.310), trimmed by trimAl (3) (1.2rev59 ; with option -automated1) and reconstructed by IQ-TREE (4, 5) (ver. 1.6.10). Two sequences from the Ascomycota phylum basal to the clade consisting of luciferase sequences were chosen as the outgroup (**Fig1D**). The sequences of five orthologues in the luciferase family—hispidin-3-hydroxylase, cytochrome P450, hispidin synthase, and caffeylpyruvate hydrolase—were constructed and the sequences were aligned by MAFFT(2) (ver. 7.310) and trimmed by trimAl(3) (1.2rev59 ; with option -automated1). The protein trees were constructed by IQ-TREE (4, 5) (ver. 1.6.10; with option -bb 10000 -alrt 1000). The protein tree of luciferase was reconciled with the species tree and the branch with < 90 SH-like bootstrap threshold were rearranged using the NOTUNG software package (21, 22). The evidence for selection across gene families was tested using the HyPhy(23, 24) platform in the webserver of datamonkey(25-27). According to the recombination breakpoints analysed by the Genetic Algorithm for Recombination Detection(28) (GARD), the alignment was trimmed for analysing selection using Single-Likelihood Ancestor Counting(29) (SLAC) and Mixed Effects Model of Evolution(30) (MEME) with *P*<0.1.

**II.** **SUPPLEMENTARY FIGURES**

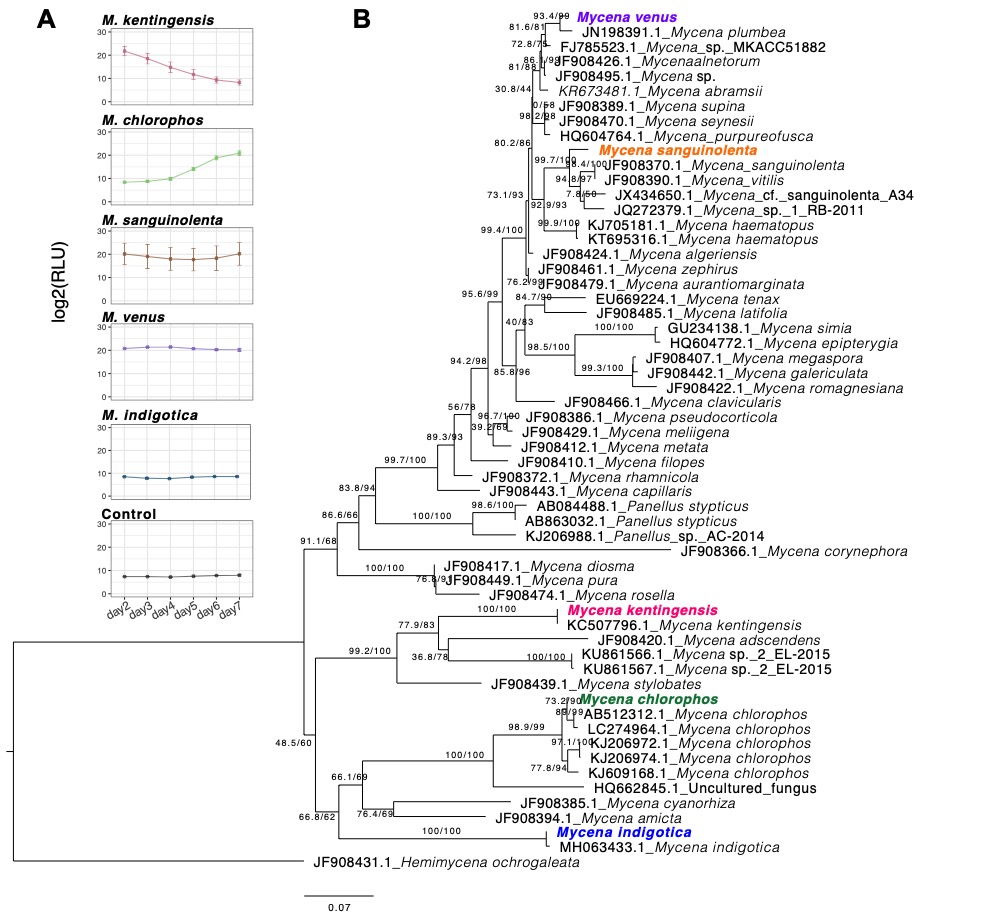

**Fig. S1. *Mycena* ITS phylogenetic tree and bioluminescent pattern in five *Mycena* species. A**, The bioluminescence patterns were measured for seven days after inoculation of mycelium on potato dextrose agar (PDA). The control was the PDA only. The diameter of mycelial growth is described in **dataset S9**. **B,** The ITS tree was constructed by IQ-TREE (4) with an alignment length of 666 bp. The isolates used in this study were denoted in colour. Numbers on each branch denote support values (SH-aLRT support (%) / ultrafast bootstrap support (%)).

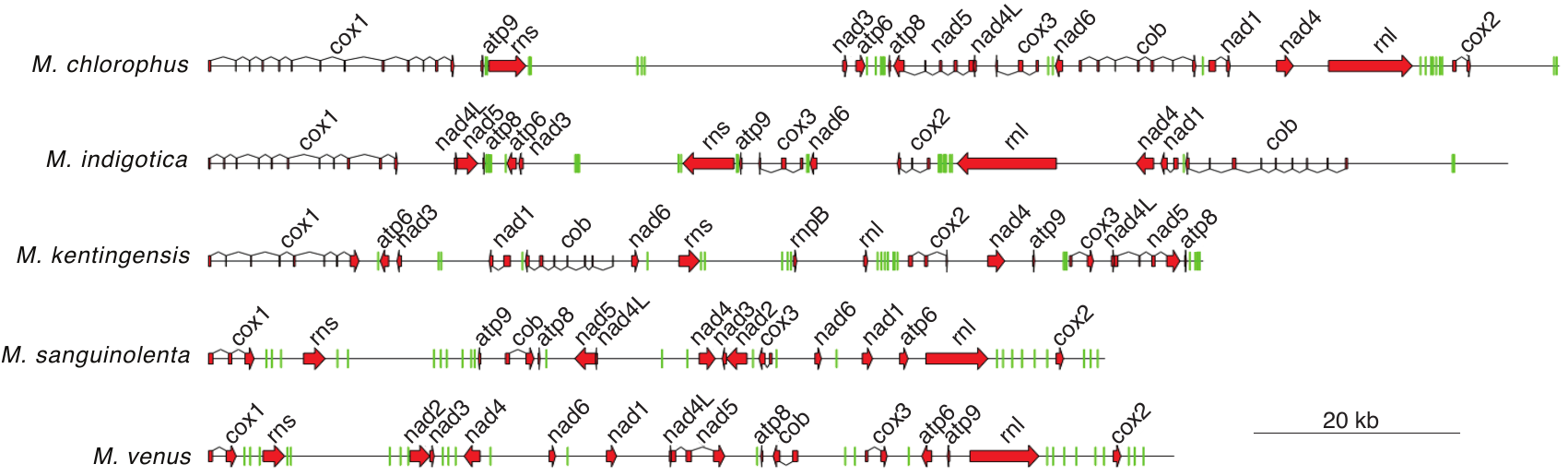

**Fig. S2. The mitochondrial genomes of five *Mycena* species.** Exons are denoted by red boxes, while introns are indicated with connecting lines. Green stripe denotes the tRNA.

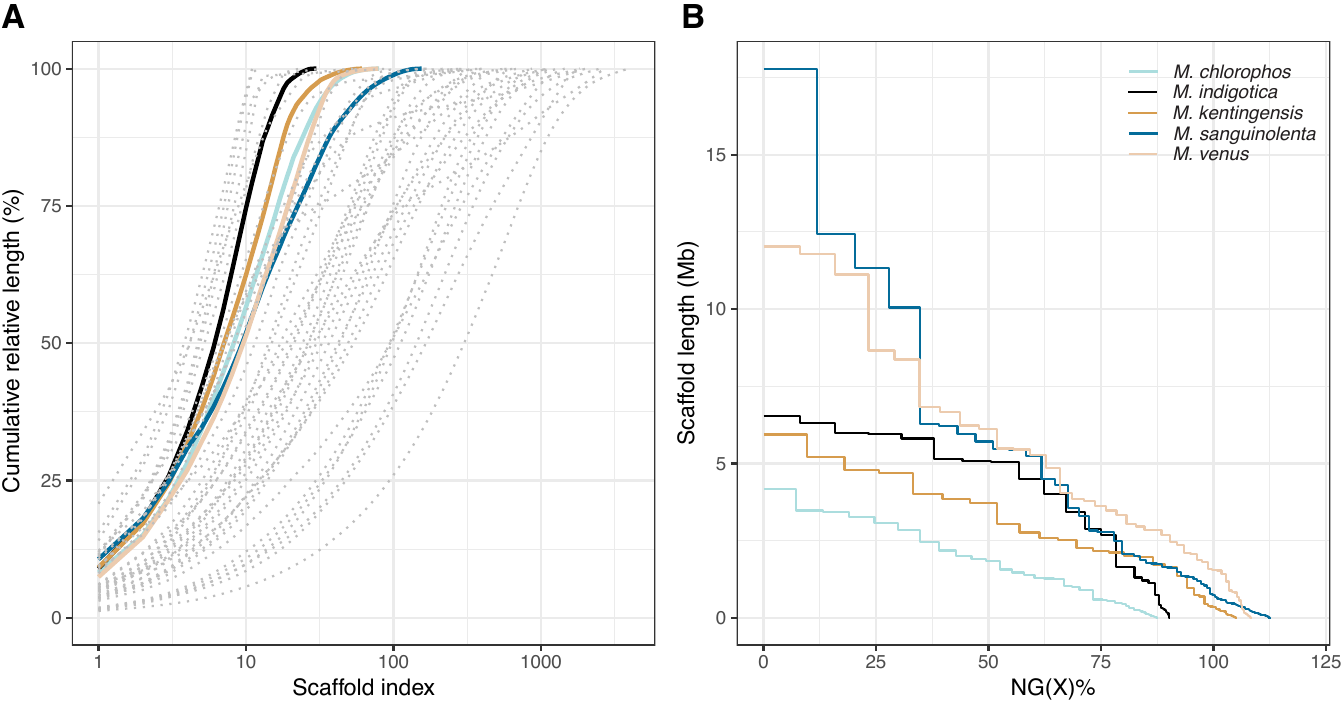

**Fig S3. (A) LG and (B) NG graphs showing an overview of *Mycena* assembly contig lengths.** Coloured and dashed lines denote *Mycena* and published assemblies used in this study, respectively. LG is the number of scaffolds sorted by length that covered cumulative percentage of the assembly. NG was calculated according to Brandnam *et al (31)* where denominator was the genome size estimated from GenomeScope (32).

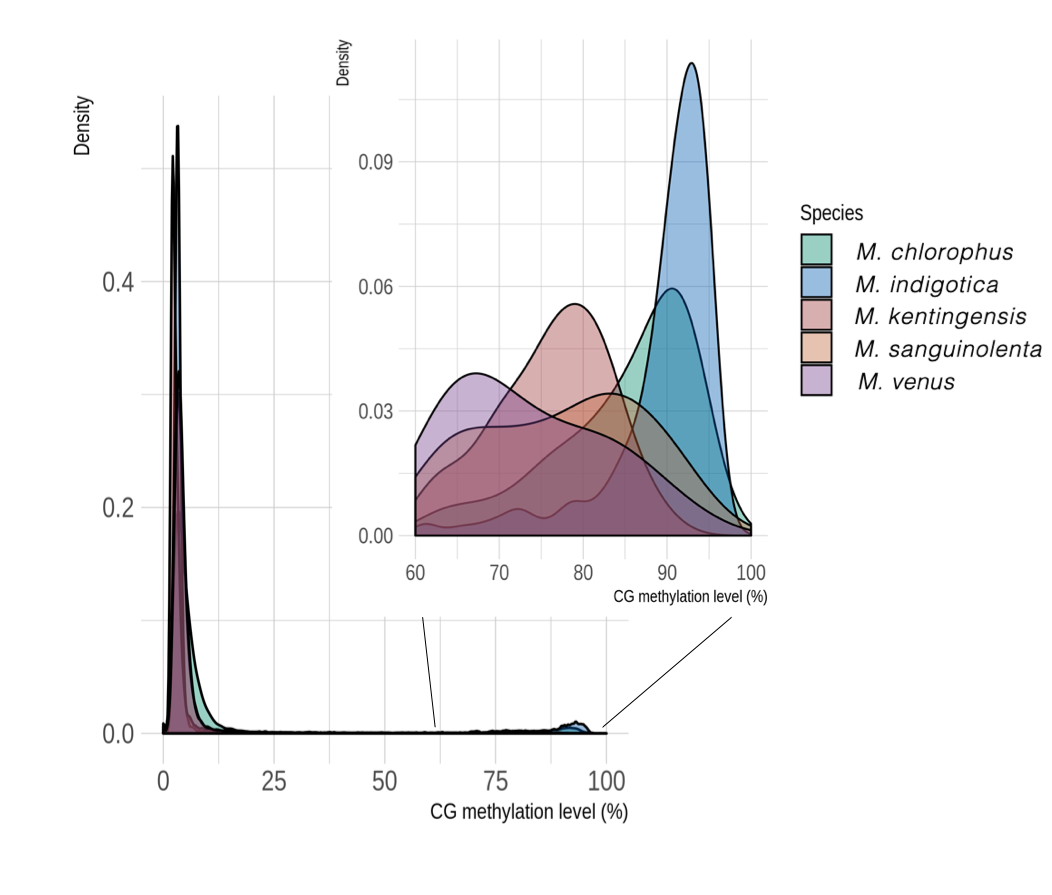

**Fig. S4.** **The distribution of CG methylation levels from gene bodies of all genes in five *Mycena* species**.

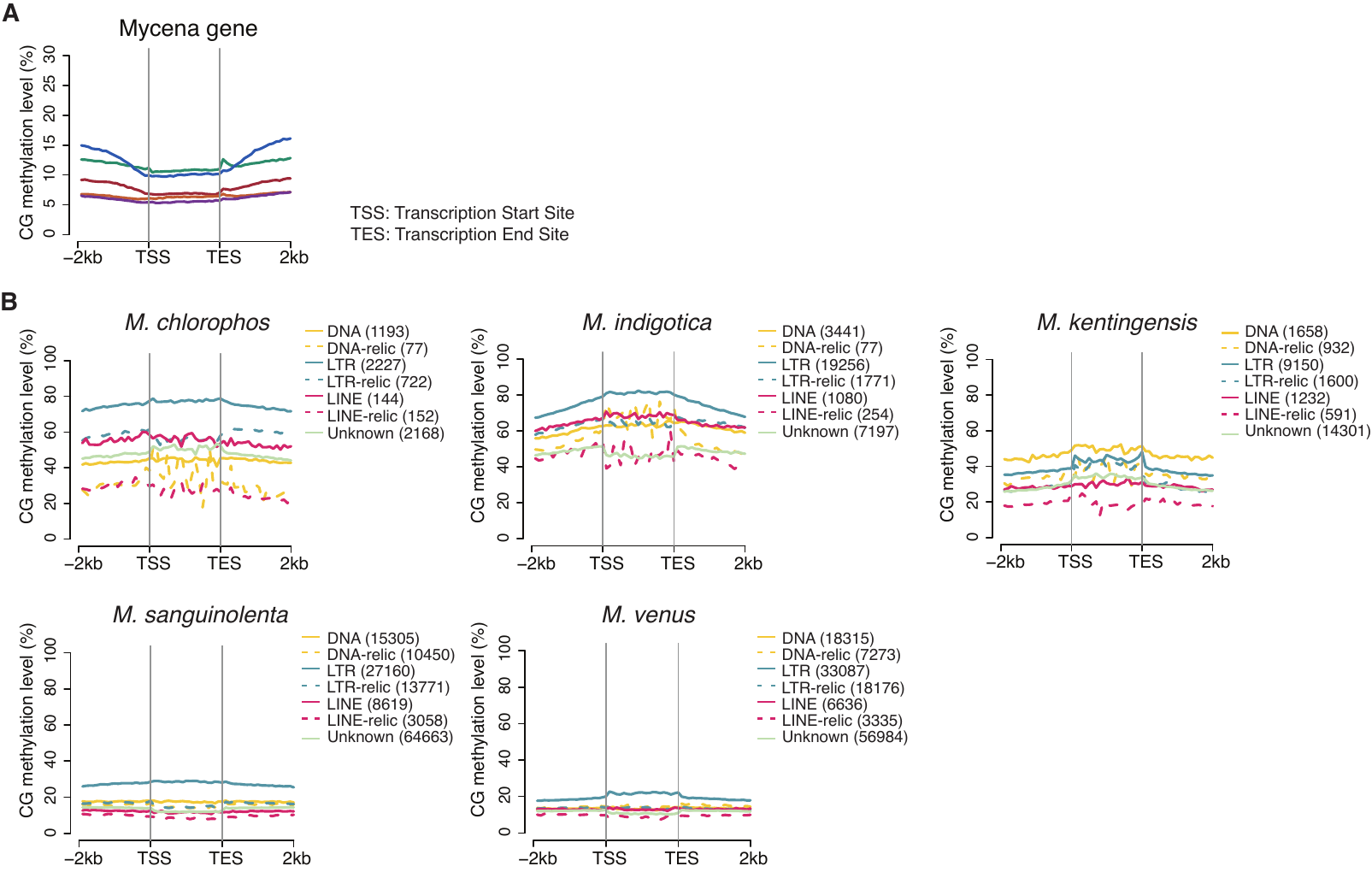

**Fig S5. CG methylation levels of (A) gene, and (B) classic TEs and their relics in 5 *Mycena* species.** Line represents a mean value of CG methylation level.

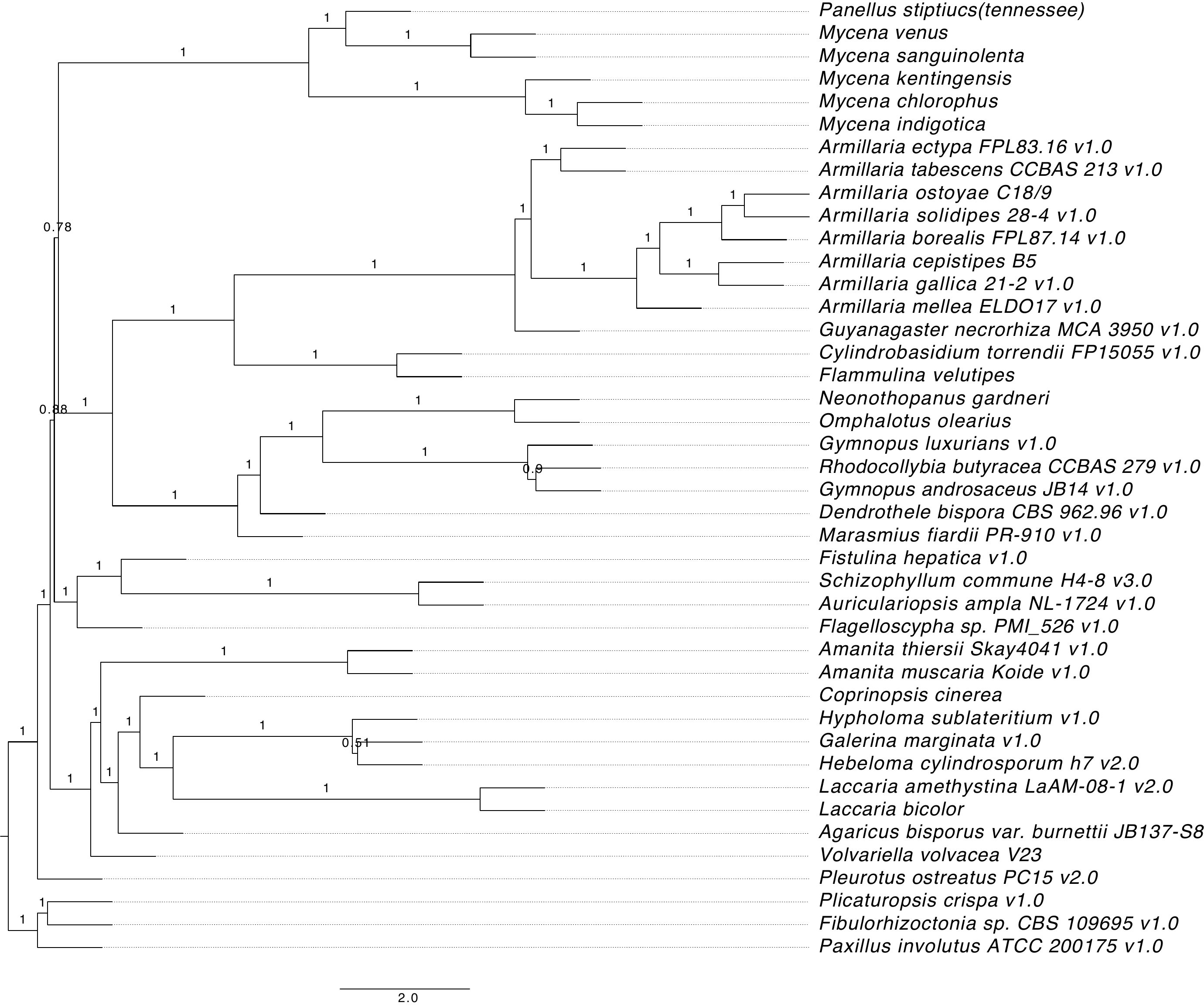

**Fig. S6. The species phylogeny reconstructed by a coalescence of 360 single-copy orthologue trees using ASTRAL-III** (33). Number on every branch represent the proportion of gene trees that support each branch

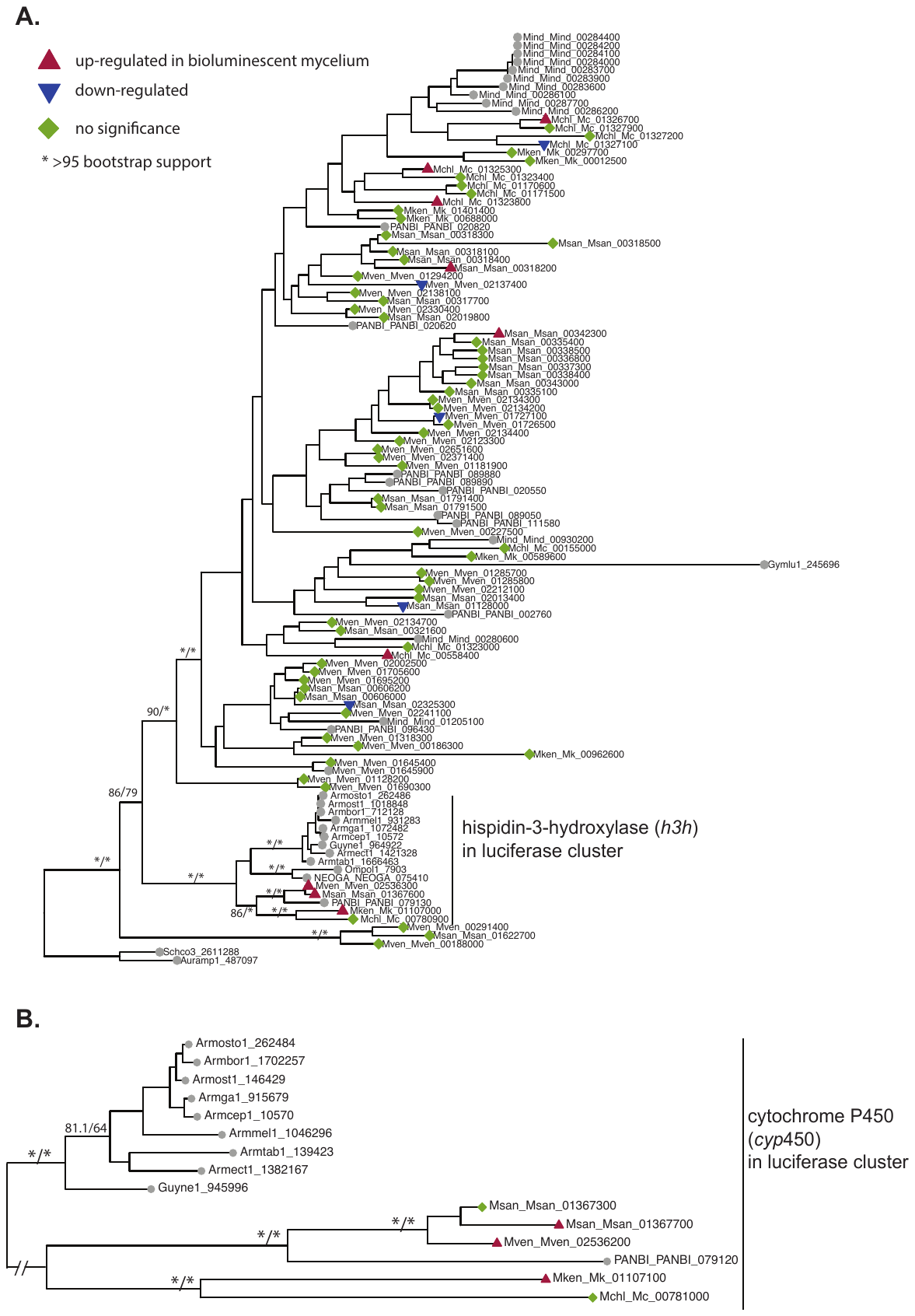

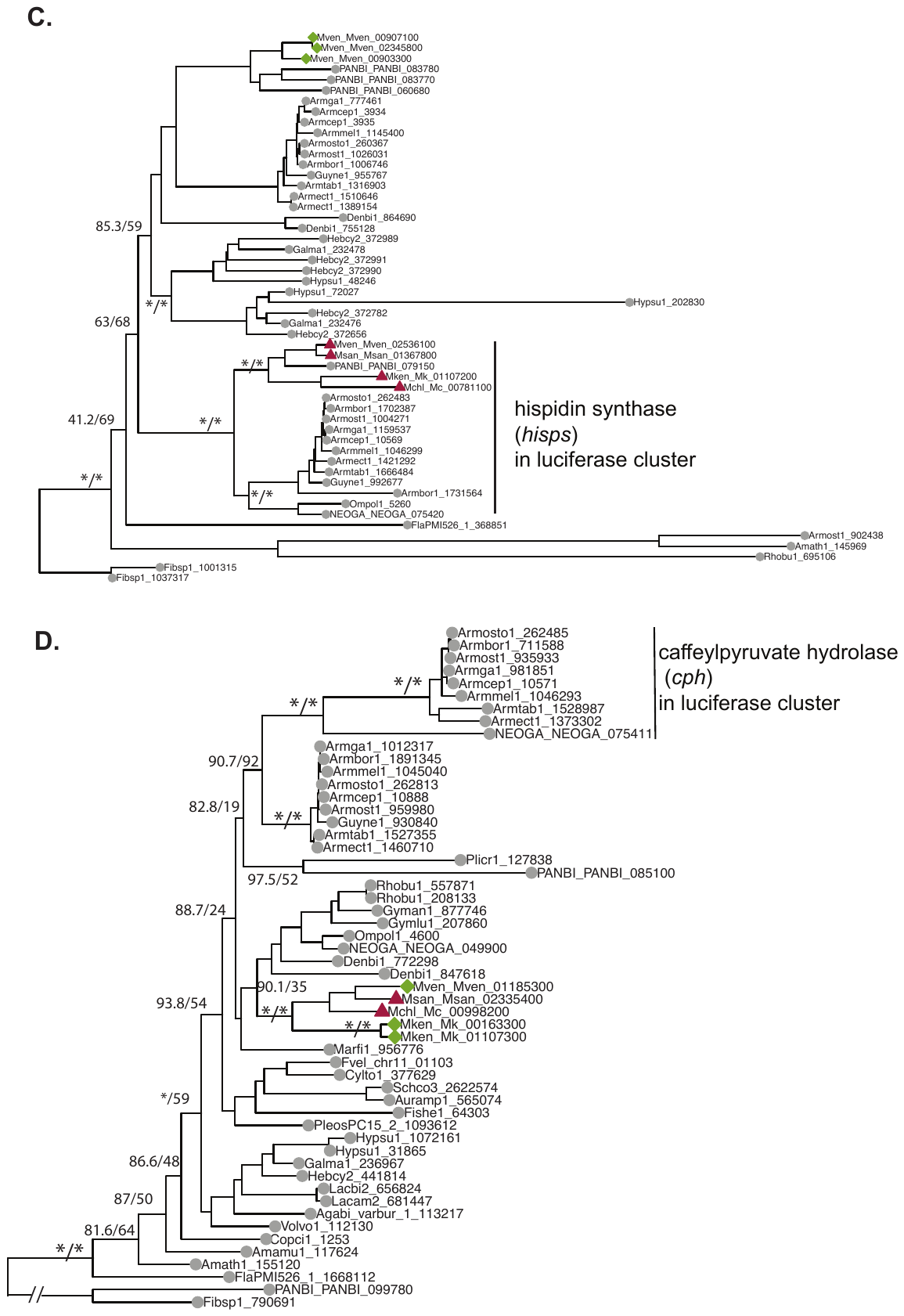

**Fig. S7.** **Phylogenies of selected gene families.** The trees were constructed by IQ-TREE(4). The value on each branch denote support values (SH-aLRT support (%) / ultrafast bootstrap support (%)) Sequence name contain species abbreviations in **Table S4** followed by gene ID. **A**, Phylogeny of OG0000706, which is the OG including hispidin-3-hydroxylase in the luciferase cluster. **B**, Phylogeny of OG0009696, which is the OG including cytochrome P450 (*cyp450*) in the *Mycena* and *Armillaria* luciferase cluster. **C,** Phylogeny of OG0002489, which is the OG including hispidin synthase (*hisps*) in the luciferase cluster. **D,** Phylogeny of OG0002332, which is the OG including caffeylpyruvate hydrolase (*cph*). For each species, the sequence id denoted the transcript ID.

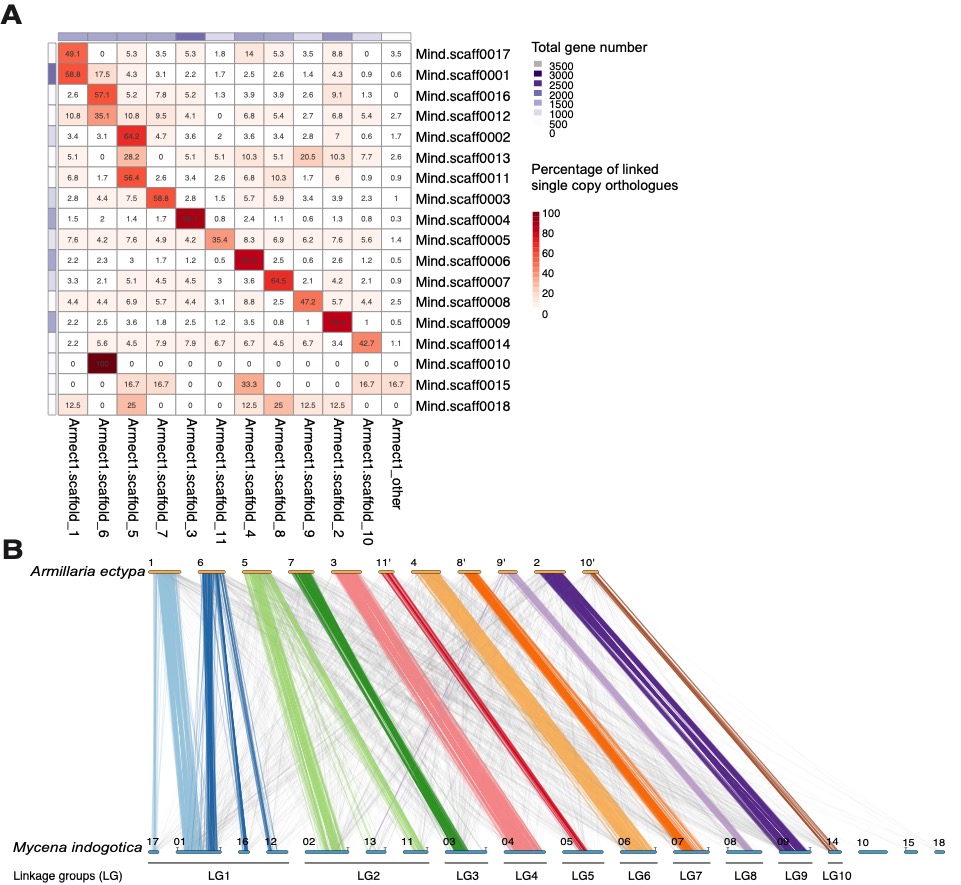

**Fig. S8**. **Synteny between *Armillaria ectypa* and *Mycena indigotica.* A**, Heatmaps of single-copy orthologues shared among scaffolds of *A. ectypa* and *M. indigotica*. Numbers in cells show the orthologues percentages of *M. indigotica* scaffold links to an *A. ectypa* scaffold. **B**, Linkage maps of single-copy orthologue linking scaffolds of *A. ectypa* and *M. indigotica*. Fifteen *M. indigotica* scaffolds were assigned unambiguously (≥ 15% of pairwise orthologues) to a corresponding *A. ectypa* scaffold, providing strong evidence that macro-synteny has been conserved across the Marasmioid clade.

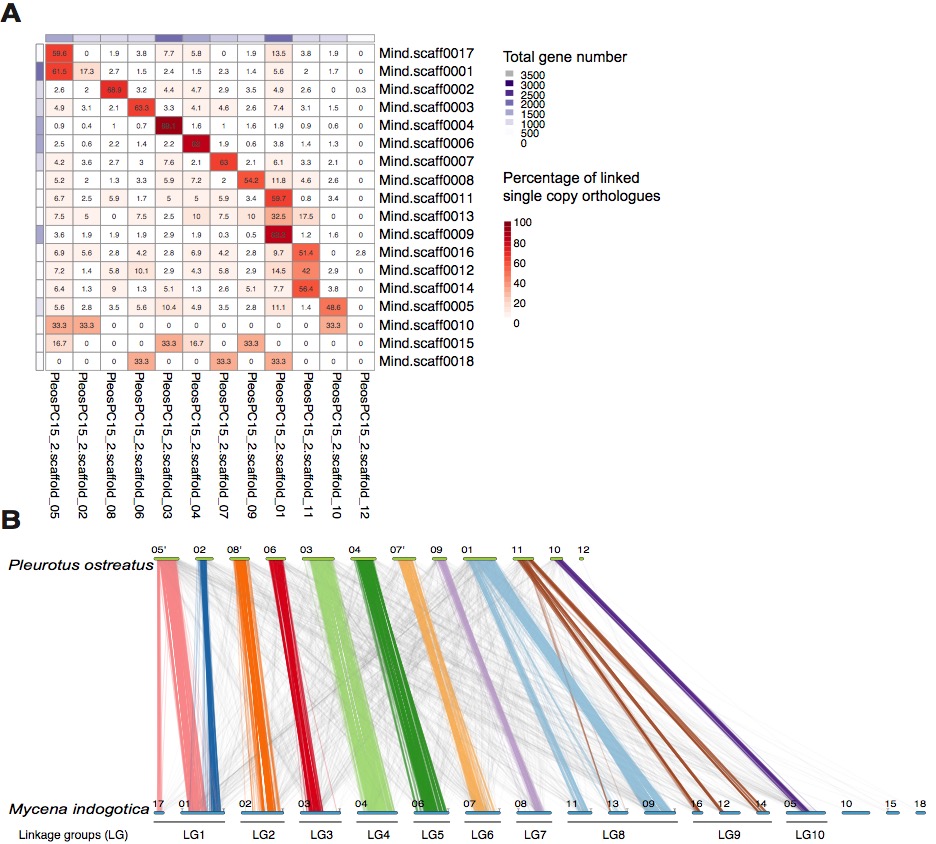

**Fig. S9.** **Synteny between *Pleurotus ostreatus* and *Mycena indigotica****.* **A**, Heatmaps of single-copy orthologues shared among scaffolds of *P. ostreatus* and *M. indigotica*. Numbers in cells show the orthologue percentages of an *M. indigotica* scaffold links to a *P. ostreatus* scaffold. **B**, Linkage maps of single-copy orthologue linking scaffolds of *P. ostreatus* and *M. indigotica*. Fifteen *M. indigotica* scaffolds were assigned unambiguously (≥ 15% of pairwise orthologues) to a corresponding *P. ostreatus* scaffold, providing strong evidence that macro-synteny has been conserved across Agaricales.

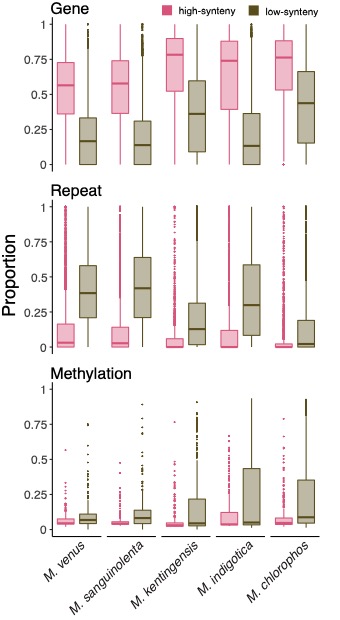

**Fig. S10.** **The gene density, repeat density and methylation level in high- or low-synteny regions.** The gene and repeat densities are calculated from the non-overlapping 10-kb window located in the core or dispensable regions. The methylation level was calculated from the mean CG methylation level in core or dispensable regions.

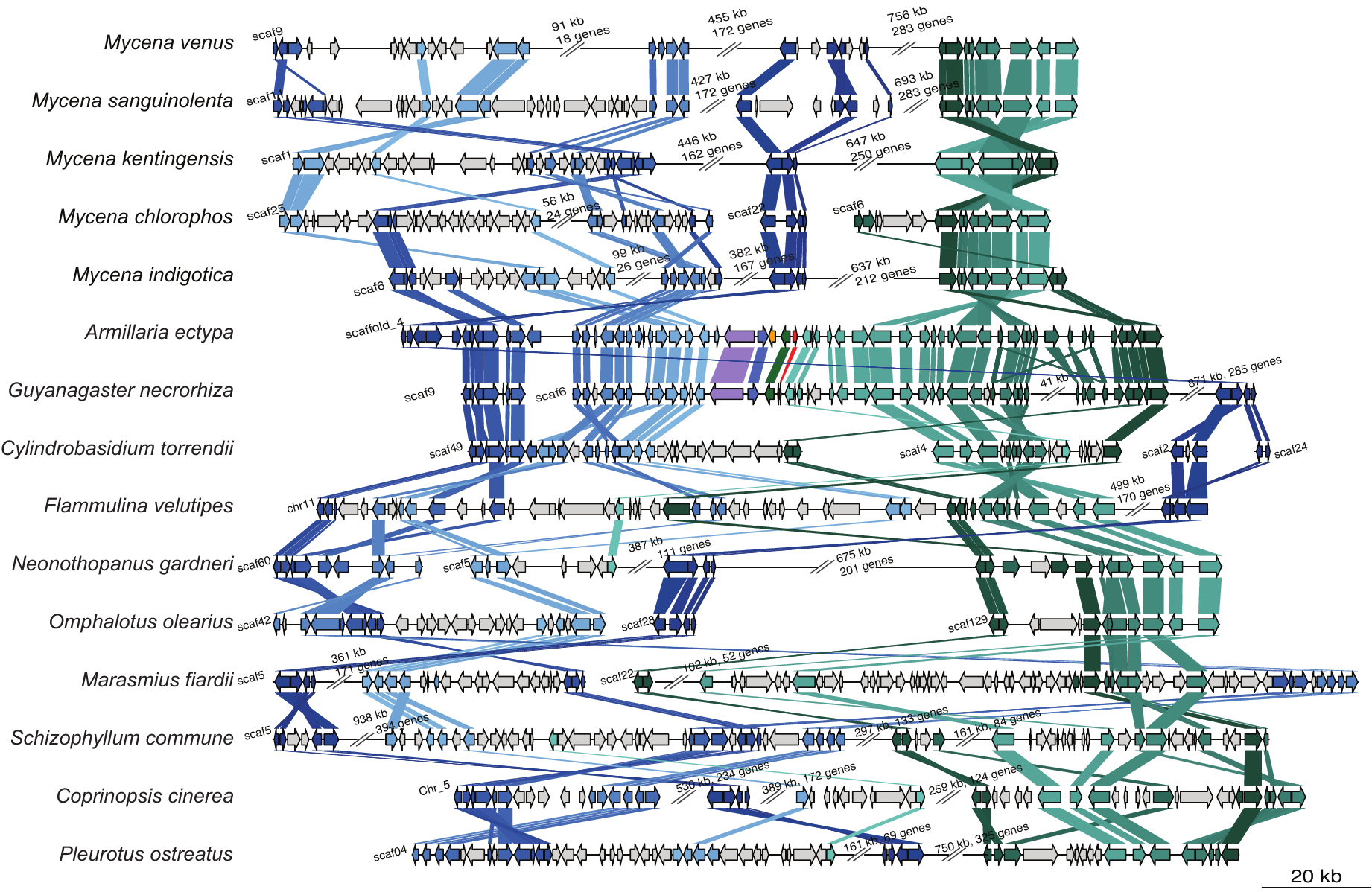

**Fig. S11. Synteny between adjacent regions of luciferase cluster in *Armillaria ectypa* and other species across Agaricales species.** Using *Armillaria ectypa* as the reference, the orthologous genes in other species are denoted with same colour regardless of their orientations. The start and stop positions for each sequence fragments: (1) *M. venus*, 4468578–5869172 of Mven.scaff0009. (2) *M. sanguinolenta*, 4419874–3166494 of Msan.scaff0011. (3) *M. kentingensis*, 3886937–2696567 of scaff0001. (4) *M. chlorophos*, 322781–451127 of Mc.scaff0025, 1443132-1451974 of Mc.scaff0022 and 904867-867954 of Mc.scaff0006. (5) *M. indigotica*, 3436952-2234518 of Mind.scaff0006. (6) *A. ectypa*, 12804877-2948226 of scaffold_4 (7) *G. necrorhiza*, 16830-4779 of scaffold_9 and 1845297-823119 of scaffold_6 (8) *C. torrendii*, 67494-130334 of scaffold_49, 345423-381212 of scaffold_4, 297084-304272 of scaffold_2, and 180513-183495 of scaffold_24. (9) *F. velutipes*, 2720572-2064978 of chr11 (10) *N. gardneri*, 2027286-2055399 of NG_scaffold_60, 28257-1173714 of NG_scaffold_5 (11) *O. olearius*, 99605-162156 of scaffold_42, 1117258-1125377 of scaffold_28, and 2738-46147 of scaffold_129 (12) *M. fiardii*, 59608-470702 of scaffold_5 and 1317903-1547407 of scaffold_22. (13) *S. commune*, 1264633-2821423 of scaffold_5. (14) *C. cinerea*, 2256009-943709 of Chr_5. (15) *P. ostreatus*, 2520611-1474541 of scaffold_04.

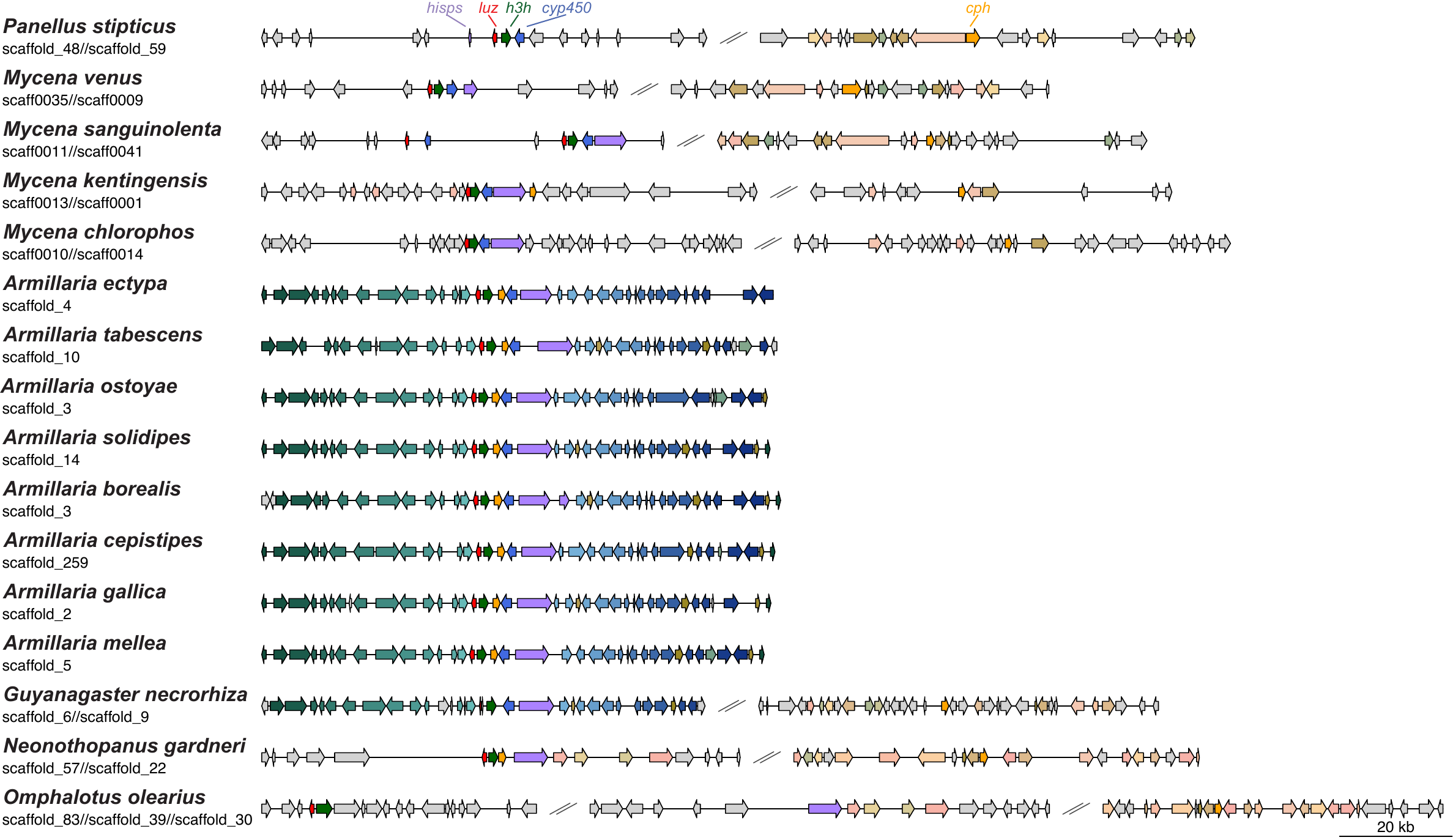

**Fig. S12.** **The similarity among genes related to bioluminescence among bioluminescent fungi and non-bioluminescent *G. necrorhiza*.** The *cph* gene in some species is locate in another scaffolds. The OGs shared by at least two species are labelled with the same colour, regardless of the orientation.

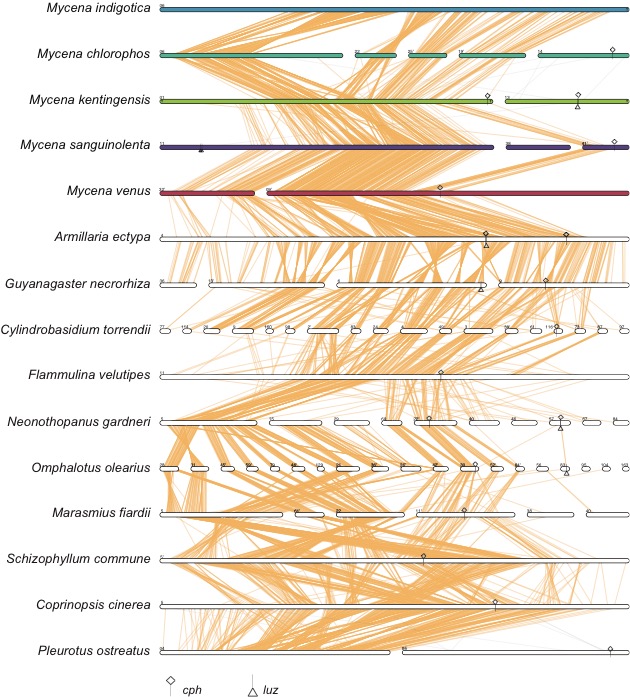

**Fig. S13. Synteny around *cph****.* Most *cph* copies (denoted by diamond) were located in different scaffolds to luciferase cluster (denoted by triangle). A copy of *cph* in *Armillaria ectypa* (Armect1) is located outside luciferase cluster shared synteny with other Agaricales suggesting this was the ancestral copy. The links denoted single-copy orthologue between two species.

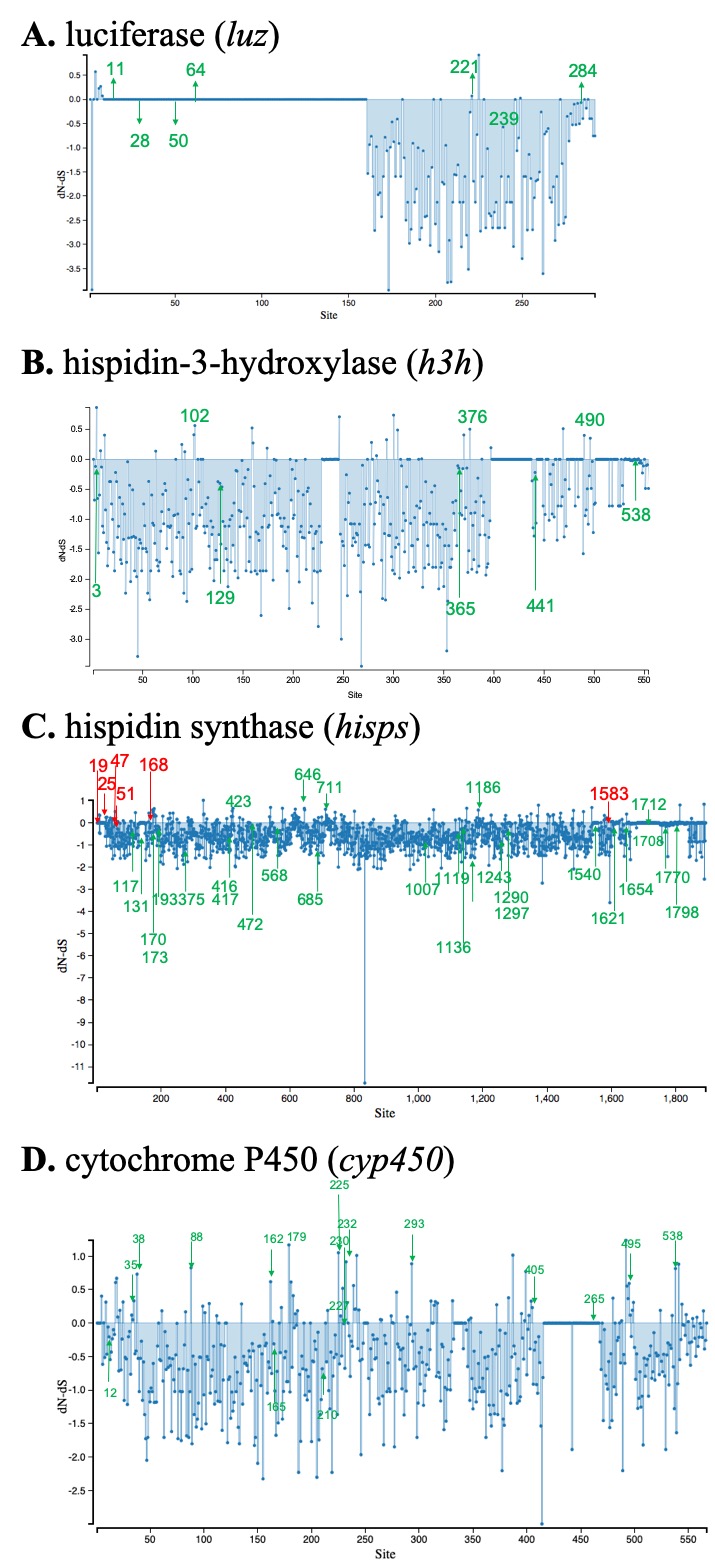

**Fig. S14. Selection analysis of the genes in luciferase cluster.**

Normalized dN/dS from SLAC (29) and MEME (30) analysis across a multispecies alignment from those in luciferase cluster. A codon position under episodic selection or positive/diversifying selection was indicated as green and red arrow, respectively. **A**, 16 *luz* sequences with 292 sites, 7 of which are under episodic selection (green arrow), and 132 of which have been under strong purifying selection. **B**, sixteen *h3h* sequences with 554 sites, eight of which are under episodic selection and 204 of which have been under strong purifying selection. **C**, seventeen *hisps* sequences with 1893 sites, twenty-eight of which are under episodic selection, six of which are under positive/diversifying selection and 637 of which have been under strong purifying selection. **D**, fifteen *cyp450* sequences with 567 sites, 17 of which are under episodic selection, and 131 of which have been under strong purifying selection.

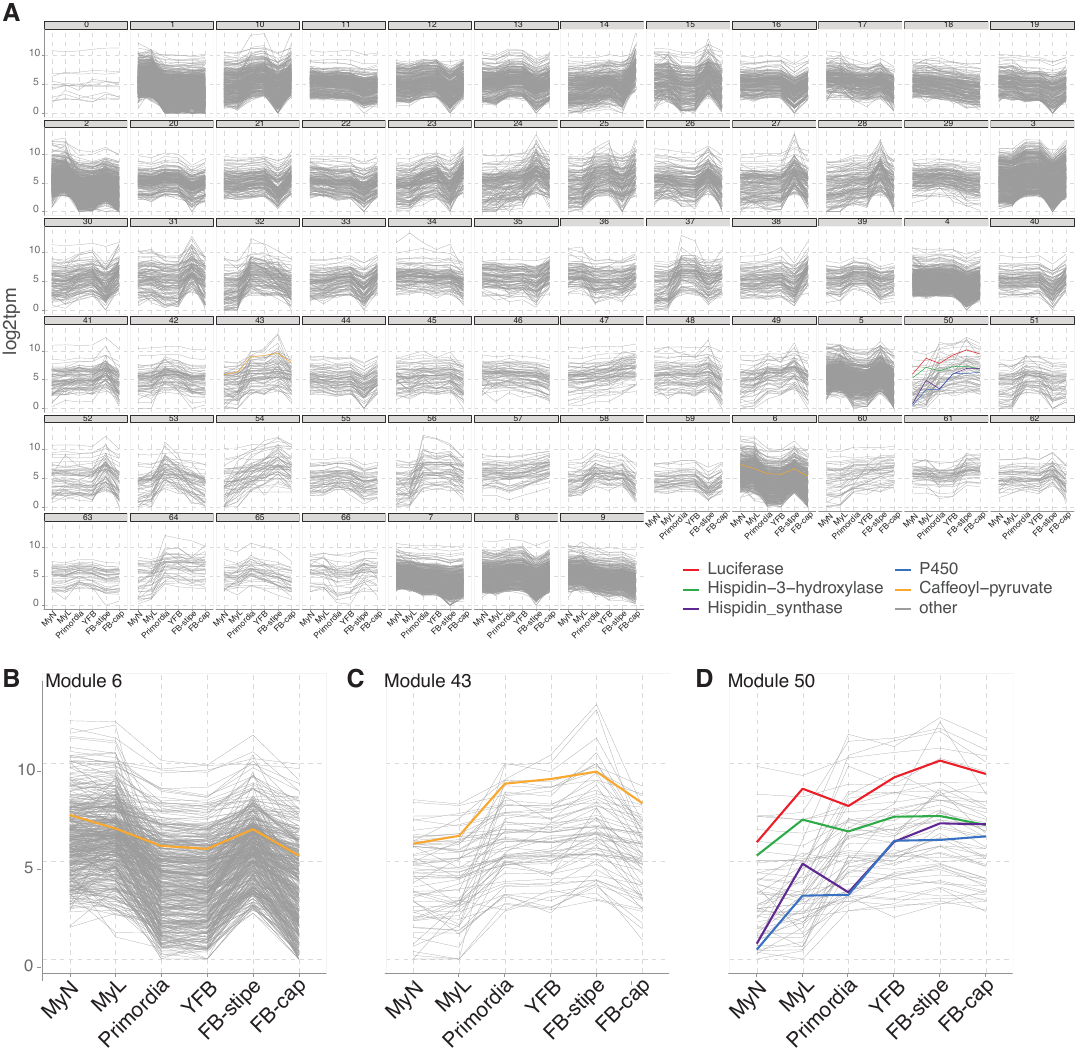

**Fig. S15.** **The 67 co-expressed gene modules identified using the weighted correlation network analysis (WGCNA) (34, 35) across developmental stages in *M. kentingensis***. **A,** All 67 modules. Different line colour denotes the expression of different genes in the luciferase cluster: red line: luciferase (*luz*); green line: hispindin-3-hydroxylase (*h3h*); blue line: cytochrome P450 (*cyp450*); purple line: hispidin synthase (*hisps*). Yellow lines are two caffeylpyruvate hydrolase (*cph*) genes assigned to different modules. **B**, One of the *cph* (Mk_00163300) located in scaffold1 was assigned to module6, whereas **C**, the other *cph* (Mk_01107300) adjacent to the luciferase cluster was assigned to module43. **D**, The genes located in luciferase cluster including *luz*, *h3h*, *cyp450*, and *hisps* were assigned into the same module, module50.

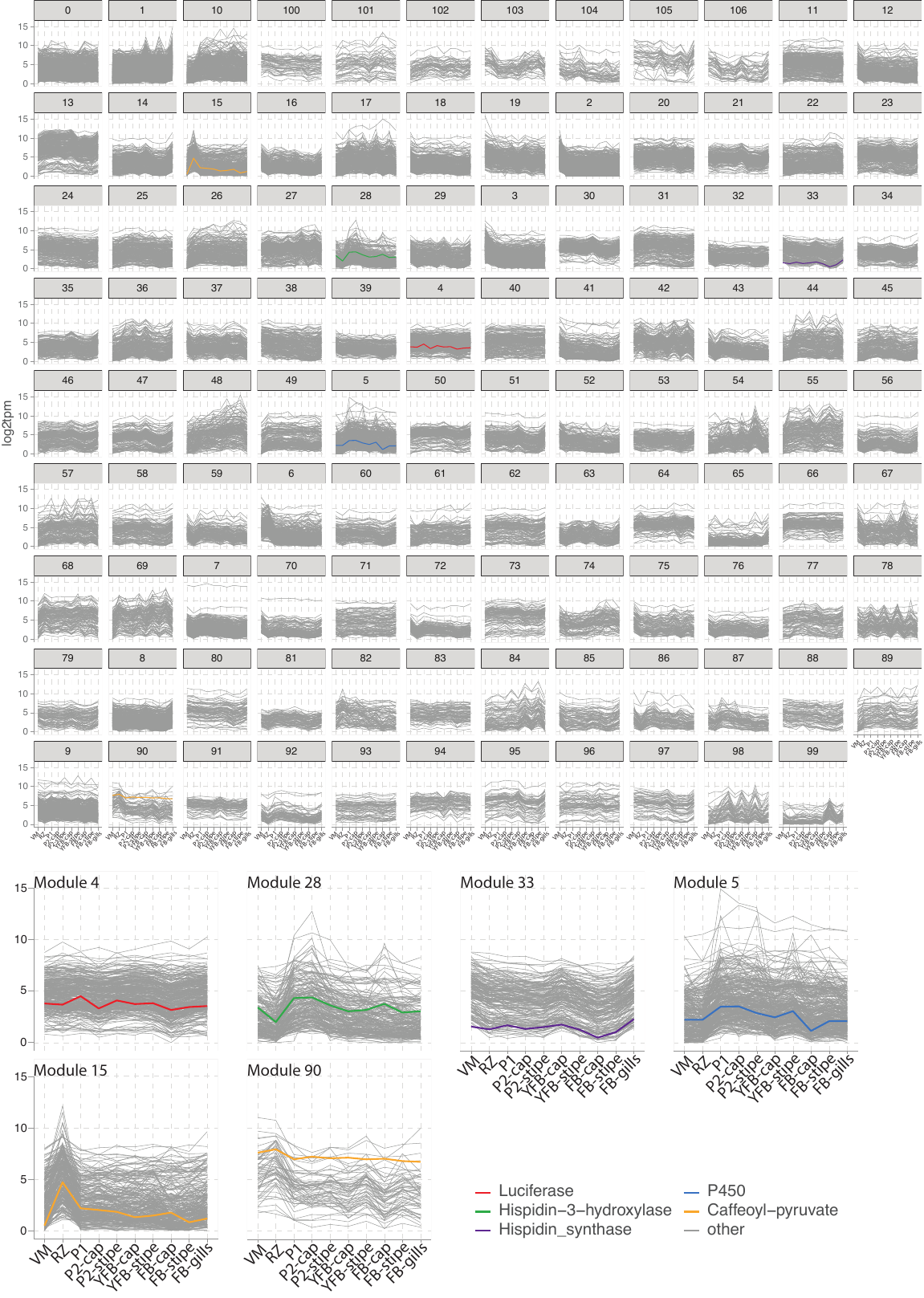

**Fig. S16.** **The 107 co-expressed gene modules identified using the weighted correlation network analysis (WGCNA)** (34, 35) **across developmental stages in *Armillaria ostoyae*.** Different line colour denotes the expression of different genes in the luciferase cluster: Red line: luciferase (*luz*); green line: hispindin-3-hydroxylase (*h3h*); blue line: cytochrome P450 (*cyp450*); yellow lines: two caffeylpyruvate hydrolase (*cph*) genes. For the two *cph*, one was located in the luciferase cluster and was assigned to Ｍodule15 and the other was located outside of the cluster and was assigned to Module90.

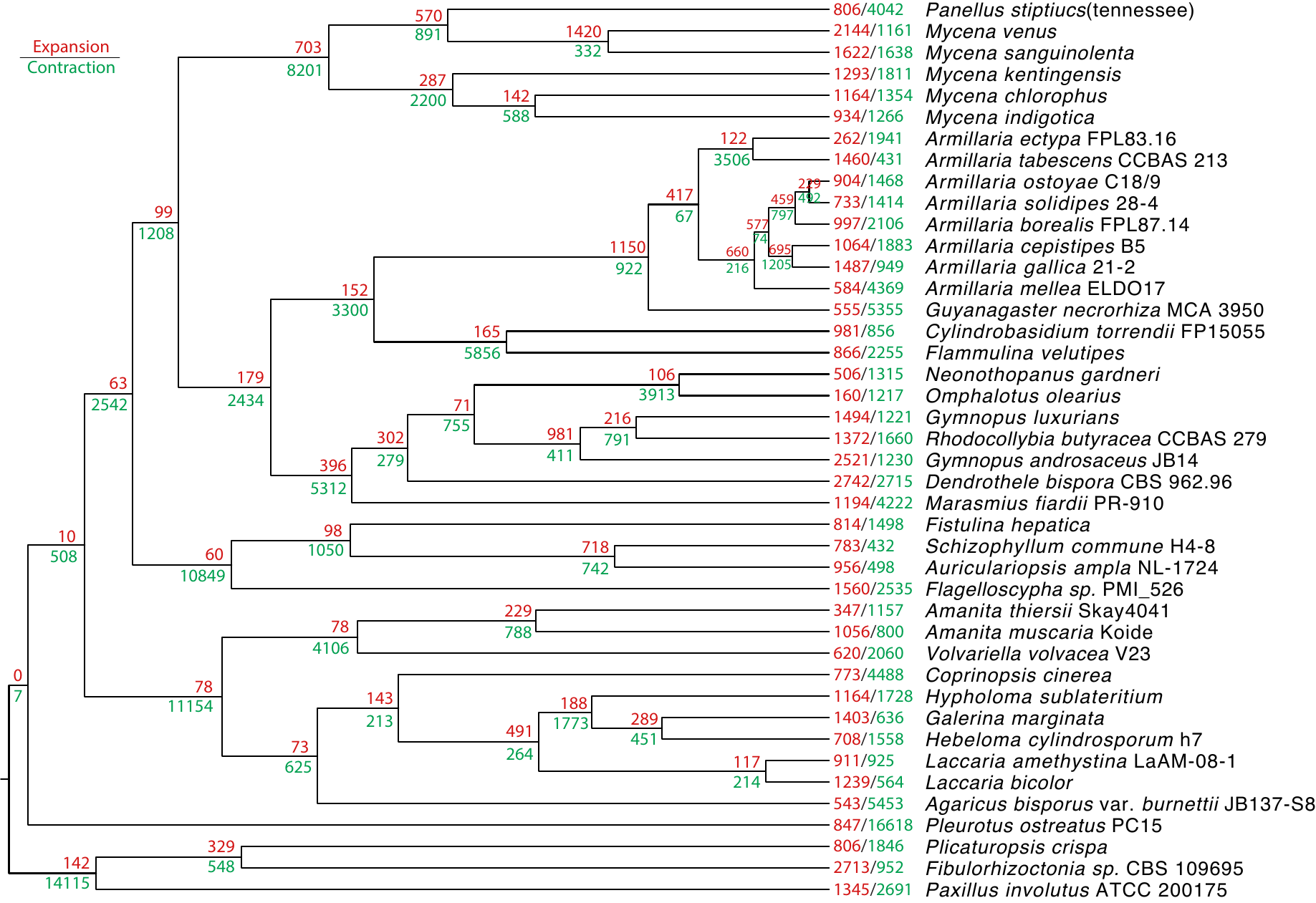

**Fig. S17. Gene expansion along the species tree of 42 basidiomycetes.** The expanded and contracted orthologous group (OG) count are coloured in red and green, respectively.

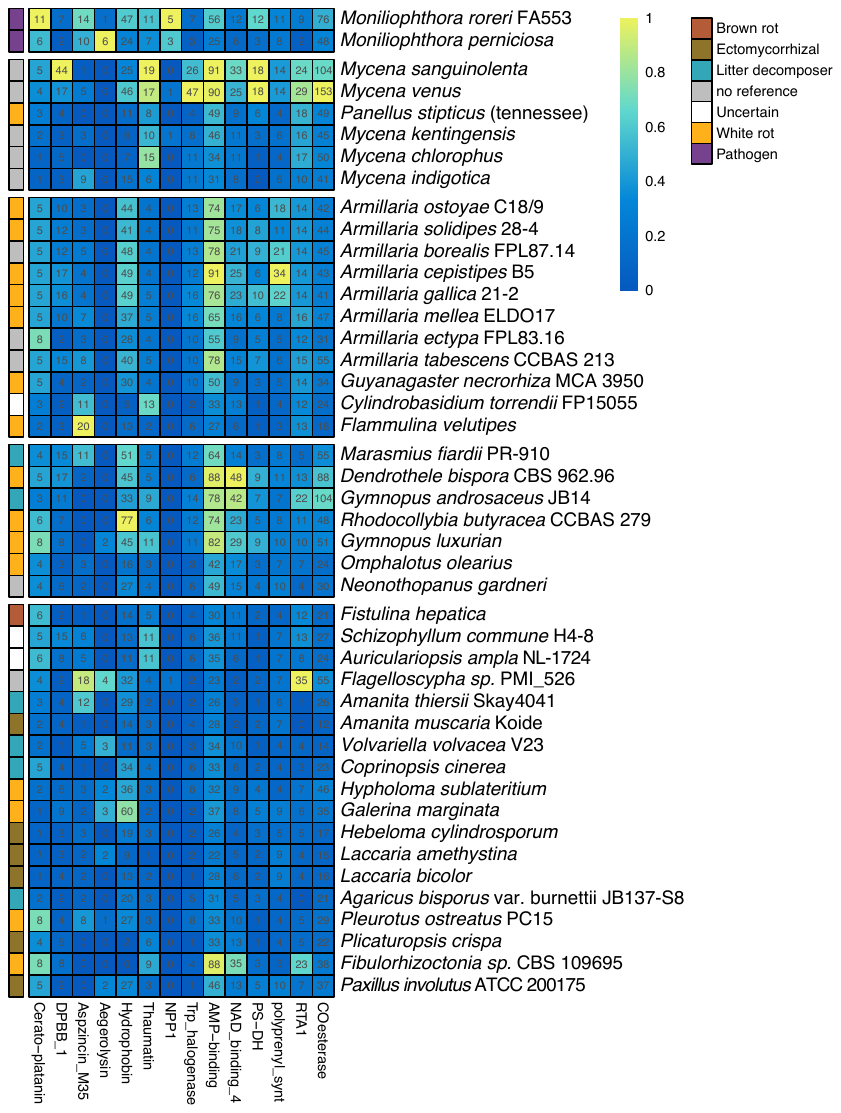

**Fig. S18. Protein family domain (Pfam) analysis across 44 fungal species.** Some Pfam domains are related to plant pathogenicity. *Moniliophthora perniciosa* FA553 (17) from JGI. Monro: *Moniliophthora roreri* from BioProject PRJNA279170. The colours from blue to yellow denote the number rescaled between 0 to 1 (for each domain, copy number from 44 species is divided by the maximum number of copies). Number inside cells denote domain copy number. The names of protein domains are labelled in the horizontal axis and the names of species are labelled in the vertical axis. Abbreviations for Pfam domain IDs and domain functions: Cerato-platanin (PF07249;Cerato-platanin), DPBB_1 (PF03330; Expansin), Aspzincin_M35 (PF14521; deuterolysin), Aegerolysin (PF06355; Hemolysin), Hydrophobin (PF01185; Hydrophobins), Thaumatin (PF00314; PR-5/Thaumatin family), NPP1 (PF05630; MpNEP1 & 2), Trp_halogenase (PF04820; Halogenase), AMP-binding (PF00501; NRPS-like synthase_1), NAD_binding_4 (PF07993; NRPS-like synthase_2), PS-DH (PF14765; Polyketide synthase), polyprenyl_synt (PF00348; Polyprenlyl Synthases), RTA1 (PF04479; RTA1_MDTM-Protein), and COesterase (PF00135; Carboxylesterase).

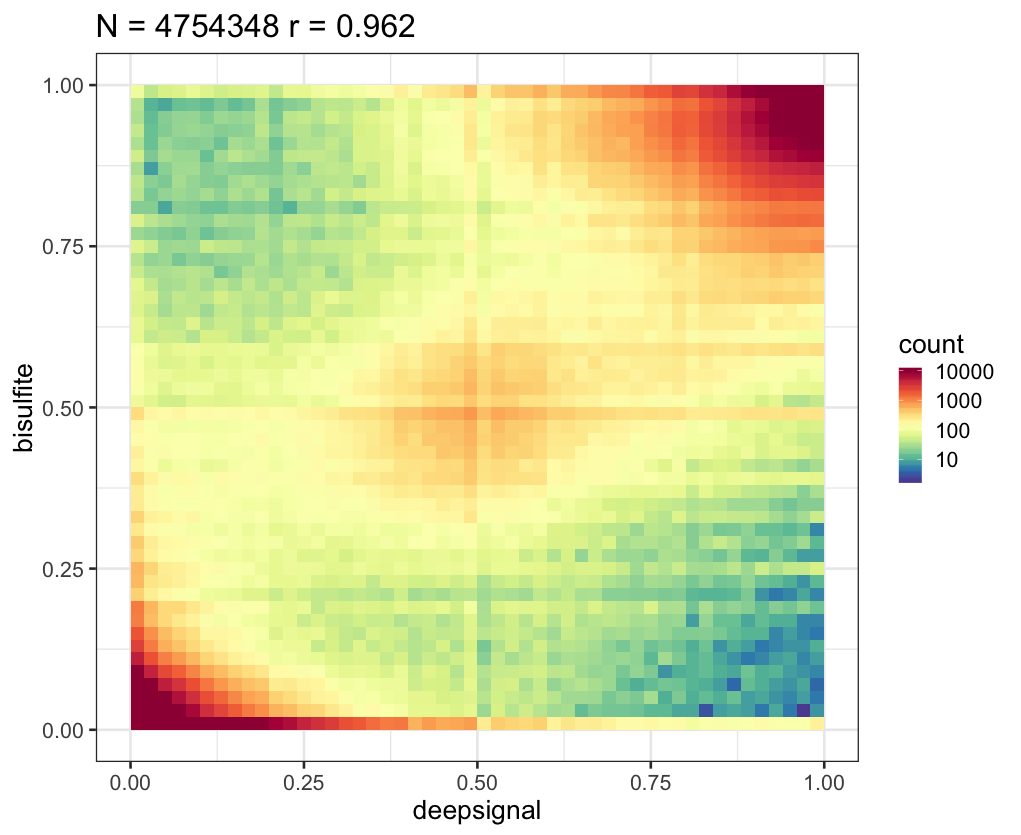

**Fig. S19. DNA methylation levels between Nanopore long-reads and the Illumina BS-seqs in the methylomes of *M. kentingensis.*** A total of 4,754,348 was compared with a Pearson correlation coefficient of 0.962.

**III. SUPPLEMENTARY TABLES**

**Table S1. Summary of Illumina and Oxford Nanopore reads.**

| **Species** | **Read type** | **Number of runs** | **Read length** | **N50** | **Library size (bp)** | **Number of reads** | **Depth of coverage** |
| --- | --- | --- | --- | --- | --- | --- | --- |
| *M. chlorophos* | Nanopore | 2 | 1,000-202,448 | 19930 | 9,375,866,055 | 965,876 | 184.3 |
| *M. indigotica* | Nanopore | 3 | 1,000-345,966 | 13971 | 15,277,834,747 | 2,240,375 | 209.6 |
| *M. kentingensis* | Nanopore | 3 | 1,000-293,280 | 26695 | 11,220,635,883 | 940,766 | 172.6 |
| *M. sanguinolenta* | Nanopore | 2 | 1,000-226,007 | 19308 | 9,755,045,881 | 1,386,206 | 58.4 |
| *M. venus* | Nanopore | 5 | 1,000-823,071 | 12272 | 21,979,383,148 | 4,187,862 | 135.5 |
| *M. chlorophos* | Illumina | 3 | 151 | N/A | 5,019,692,396 | 33,242,996 | 98.7 |
| *M. indigotica* | Illumina | 1 | 151 | N/A | 5,957,541,014 | 39,453,914 | 81.7 |
| *M. kentingensis* | Illumina | 3 | 151, 301 | N/A | 4,118,183,612 | 21,304,412 | 63.3 |
| *M. sanguinolenta* | Illumina | 4 | 151 | N/A | 18,276,688,774 | 121,037,674 | 109.3 |
| *M. venus* | Illumina | 3 | 151 | N/A | 18,266,335,610 | 120,969,110 | 112.6 |

**Table S2. Genome summaries of five *Mycena* species.**

Genome haploid length and heterozygosity of each species was calculated from Illumina reads using GenomeScope 2.0 (32)

|  | ***M. chlorophos*** | ***M. indigotica*** | ***M. kentingensis*** | ***M. sanguinolenta*** | ***M. venus*** |
| --- | --- | --- | --- | --- | --- |
| Abbreviation in this study | Mchl | Mind | Mken | Msan | Mven |
| Genome haploid length estimate (Mb) | 58.2 | 80.8 | 62.0 | 148.7 | 149.9 |
| Canu (36) assembly size (Mb) | 75.4 | 73.5 | 97.1 | 263.7 | 265.4 |
| Haplotype aware assembly size (Mb) | 50.9 | 72.9 | 65.0 | 167.2 | 162.2 |
| Estimated heterozygosity (%) | 2.92 | 0.03 | 1.7 | 2.71 | 2.35 |
| Genome size (Mb) | 50.9 | 72.9 | 65.0 | 167.2 | 162.2 |
| scaffold number (n) | 80 | 30 | 61 | 155 | 79 |
| N50 (Mb) | 2.0 | 5.1 | 3.0 | 5.4 | 5.5 |
| L50 | 9 | 7 | 8 | 10 | 10 |
| N90 (Mb) | 478,732 | 1,639,376 | 968,399 | 608,068 | 1,860,701 |
| L90 | 27 | 15 | 20 | 43 | 29 |
| Quality Value (QV) | 31.1 | 36.3 | 36.8 | 34.0 | 34.2 |
| Number of genes | 14,274 | 14,334 | 15,046 | 25,350 | 26,518 |
| Gene length (Mb) | 26.4 | 24.5 | 28.5 | 44.0 | 46.0 |
| Exon number | 89,139 | 90,123 | 96,716 | 169,212 | 181,086 |
| Exon length (Mb) | 20.3 | 20.1 | 21.8 | 33.5 | 35.2 |
| Intergene length (Mb) | 24.5 | 48.4 | 36.5 | 123.1 | 116.2 |

**Table S3.** **BUSCO summaries of five *Mycena* species based on 1,764 basidiomycete markers.**

|  | **Complete** | **Single** | **Duplicate** | **Fragmented** | **Missing** |
| --- | --- | --- | --- | --- | --- |
| *M. venus* | 92.1% | 86.1% | 6.0% | 2.0% | 5.9% |
| *M. sanguinolenta* | 92.8% | 88.1% | 4.7% | 2.0% | 5.2% |
| *M. kentingensis* | 94.2% | 89.2% | 5.0% | 1.0% | 4.8% |
| *M. chlorophos* | 95.3% | 89.1% | 6.2% | 0.7% | 4.0% |
| *M. indigotica* | 94.8% | 92.7% | 2.1% | 0.6% | 4.6% |

**Table S4.** **Summaries of 37 representative fungal species used in this study**

| **Species abbreviation** | **Species** | **Assembly**  **version** | **Reference** | **Assembly size (bp)** | **Number of**  **scaffolds** |
| --- | --- | --- | --- | --- | --- |
| Armosto1 | *Armillaria ostoyae* C18/9 |  | JGI; Sipos *et al*. (37) | 60,106,801 | 106 |
| Armost1 | *Armillaria solidipes* 28-4 | v1.0 | JGI; Sipos *et al*. (37) | 58,009,494 | 229 |
| Armbor1 | *Armillaria borealis* FPL87.14 | v1.0 | JGI with permission | 71,689,880 | 864 |
| Armcep1 | *Armillaria cepistipes* B5 |  | JGI; Sipos *et al*. (37) | 75,828,441 | 287 |
| Armga1 | *Armillaria gallica* 21-2 | v1.0 | JGI; Sipos *et al*. (37) | 85,336,812 | 319 |
| Armmel1 | *Armillaria mellea* ELDO17 | v1.0 | JGI with permission | 70,856,304 | 474 |
| Armect1 | *Armillaria ectypa* FPL83.16 | v1.0 | JGI with permission | 40,598,130 | 33 |
| Armtab1 | *Armillaria tabescens* CCBAS 213 | v1.0 | JGI with permission | 74,875,987 | 659 |
| Guyne1 | *Guyanagaster necrorhiza* MCA 3950 | v1.0 | JGI with permission | 53,686,691 | 168 |
| Cylto1 | *Cylindrobasidium torrendii* FP15055 | v1.0 | JGI; Floudas *et al*. (38) | 31,574,086 | 1,149 |
| Fvel | *Flammulina velutipes* |  | JGI with permission | 35,642,541 | 11 |
| Rhobu1 | *Rhodocollybia butyrace*a CCBAS 279 | v1.0 | JGI with permission | 78,814,333 | 1,836 |
| Gymlu1 | *Gymnopus luxurians* | v1.0 | JGI; Kohler *et al*. (39) | 66,281,680 | 383 |
| Gyman1 | *Gymnopus androsaceus* JB14 | v1.0 | JGI; Barbi *et al*. (40) | 89,149,538 | 2,516 |
| Ompol1 | *Omphalotus olearius* |  | JGI; Wawrzyn *et al*. (41) | 28,145,093 | 868 |
| NEOGA | *Neonothopanus gardneri* |  | Kotlobay *et al*. (42) | 46,745,715 | 293 |
| Denbi1 | *Dendrothele bispora* CBS 962.96 | v1.0 | JGI; Varga *et al*. (43) | 130,650,616 | 3,942 |
| Marfi1 | *Marasmius fiardii* PR-910 | v1.0 | JGI with permission | 59,447,912 | 1,124 |
| PANBI | *Panellus stiptiucs*(tennessee) |  | Kotlobay *et al*. (42) | 39,545,155 | 571 |
| Schco3 | *Schizophyllum commune* H4-8 | v3.0 | JGI; Ohm *et al* (44) | 38,670,379 | 25 |
| Auramp1 | *Auriculariopsis ampla* NL-1724 | v1.0 | JGI; Almasi *et al*. (45) | 49,873,921 | 351 |
| Fishe1 | *Fistulina hepatica* | v1.0 | JGI; Floudas *et al*. (38) | 33,847,808 | 588 |
| FlaPMI526_1 | *Flagelloscypha sp.* PMI_526 | v1.0 | JGI with permission | 72,728,845 | 397 |
| Galma1 | *Galerina marginata* | v1.0 | JGI; Riley *et al*. (46) | 59,418,196 | 414 |
| Hebcy2 | *Hebeloma cylindrosporum* h7 | v2.0 | JGI; Kohler *et al*. (39) | 38,226,047 | 176 |
| Hypsu1 | *Hypholoma sublateritium* | v1.0 | JGI; Kohler *et al*. (39) | 48,031,814 | 704 |
| Lacam2 | *Laccaria amethystina* LaAM-08-1 | v2.0 | JGI; Kohler *et al*. (39) | 52,581,404 | 1,299 |
| Lacbi2 | *Laccaria bicolor* |  | JGI; Kohler *et al*. (39) | 60,707,050 | 55 |
| Copci1 | *Coprinopsis cinerea* |  | JGI; Stajich *et al*. (47) | 36,294,355 | 94 |
| Agabi_varbur_1 | *Agaricus bisporus* var. burnettii JB137-S8 |  | JGI; Morin *et al*. (48) | 32,614,401 | 2,016 |
| Amath1 | *Amanita thiersii* Skay4041 | v1.0 | JGI; Hess *et al*. (49) | 33,689,220 | 1,446 |
| Amamu1 | *Amanita muscaria* Koide | v1.0 | JGI; Kohler *et al.* (39) | 40,699,759 | 1,101 |
| Volvo1 | *Volvariella volvacea* | v23 | JGI; Bao *et al*. (50) | 35,720,033 | 62 |
| PleosPC15_2 | *Pleurotus ostreatus* PC15 | v2.0 | JGI; Riley *et al*. (46) | 34,343,005 | 12 |
| Plicr1 | *Plicaturopsis crispa* | v1.0 | JGI; Kohler *et al*. (39) | 34,498,416 | 316 |
| Fibsp1 | *Fibulorhizoctonia sp.* CBS 109695 | v1.0 | JGI; Nagy *et al*. (51) | 95,125,689 | 1,918 |
| Paxin1 | *Paxillus involutus* ATCC 200175 | v1.0 | JGI; Kohler *et al*. (39) | 58,301,126 | 2,681 |

**Table S5. Statistics of orthologous groups (OGs) of five *Mycena* species.** A total of 22,244 OGs were inferred in 42 species.

| **Species** | **Number of OGs** | **Species-specific OGs** | **Singleton** |
| --- | --- | --- | --- |
| *M. chlorophos* | 9048 | 1378 | 1357 |
| *M. indigotica* | 9055 | 1329 | 1310 |
| *M. kentingensis* | 9295 | 1587 | 1560 |
| *M. sanguinolenta* | 12345 | 3588 | 3547 |
| *M. venus* | 12471 | 3374 | 3348 |

**Table S6. Intron length (bp) of intron-containing genes in five *Mycena* mitogenomes.** Brackets denote of number of introns. Seven genes (*nad3, nad4, nad4L, nad6, atp6, atp8* and *atp9*) are intron free.

|  | ***M. chlorophos*** | ***M. indigotica*** | ***M. kentingensis*** | ***M. sanguinolenta*** | ***M. venus*** |
| --- | --- | --- | --- | --- | --- |
| rns | 1,632 (1) | 2,923 (2) | 0 (0) | 0 (0) | 0 (0) |
| rnl | 4,611 (5) | 5,817 (5) | 3,584 (4) | 0 (0) | 0 (0) |
| cob | 10,295 (6) | 14,753 (8) | 7,632 (6) | 1,627 (1) | 1,259 (1) |
| cox1 | 22,578 (12) | 17,250 (10) | 13,223 (7) | 2,832 (2) | 1,348 (1) |
| cox2 | 1,088 (1) | 2,546 (2) | 3,050 (2) | 0 (0) | 0 (0) |
| cox3 | 3,397 (2) | 3,512 (2) | 1,531 (1) | 423 (1) | 1,295 (1) |
| nad1 | 1,093 (1) | 639 (1) | 1,079 (1) | 0 (0) | 0 (0) |
| nad2 | 5,116 (3) | 2,892 (2) | 1,563 (1) | 0 (0) | 0 (0) |
| nad5 | 5,799 (4) | 0 (0) | 4,426 (3) | 0 (0) | 3,223 (2) |
| sum | 55,609 (35) | 50,332 (32) | 36,088 (25) | 4,882 (4) | 7,125 (5) |

**Table S7. Repeat content of five *Mycena* genomes**

| **Species name** | **SINES (%)** | **LINES (%)** | **LTRs (%)** | **DNA (%)** | **Unclassified (%)** | **Small (%)** | **Satellite (%)** | **Simple (%)** | **Low (%)** | **Sum (%)** |
| --- | --- | --- | --- | --- | --- | --- | --- | --- | --- | --- |
| *M. chlorophos* | 0.14 | 0.29 | 5.22 | 1.4 | 3.97 | 0 | 0 | 0.62 | 0.08 | 11.72 |
| *M. kentingensis* | 0 | 1.06 | 6.54 | 1.53 | 8.33 | 0 | 0.04 | 0.67 | 0.1 | 18.27 |
| *M. indigotica* | 0 | 0.99 | 19.71 | 2.63 | 5.28 | 0.37 | 0 | 0.43 | 0.07 | 29.48 |
| *M. sanguinolenta* | 0.7 | 1.99 | 9.69 | 3.91 | 21.58 | 0.16 | 0.08 | 0.8 | 0.07 | 38.98 |
| *M. venus* | 0 | 1.7 | 11.09 | 4.01 | 17.94 | 0.09 | 0.06 | 0.75 | 0.06 | 35.7 |

**Table S8. Unknown and relic repeat content (%) of genomes.**

|  | **Unknown (%)** | **LTR relic**  **(%)** | **LINE relic**  **(%)** | **DNA relic**  **(%)** | **All Relic**  **(%)** |
| --- | --- | --- | --- | --- | --- |
| *M. chlorophos* | 2.73 | 0.91 | 0.16 | 0.04 | 1.11 |
| *M. kentingensis* | 6.85 | 0.70 | 0.36 | 0.32 | 1.38 |
| *M. indigotica* | 3.96 | 0.99 | 0.06 | 0.28 | 1.34 |
| *M. sanguinolenta* | 13.56 | 4.33 | 0.93 | 2.13 | 7.38 |
| *M. venus* | 10.46 | 4.62 | 0.56 | 1.37 | 6.55 |

**Table S9. CG methylation in genes, TEs, relic TEs, and unknown repeats.**

|  | ***M. chlorophos*** | ***M. indigotica*** | ***M. kentingensis*** | ***M. sanguinolenta*** | ***M. venus*** |
| --- | --- | --- | --- | --- | --- |
| Gene (%) | 10.5 | 10.1 | 6.6 | 6.2 | 5.4 |
| DNA (%) | 43.3 | 57.9 | 45.4 | 17.0 | 13.1 |
| relic DNA (%) | 34.1 | 41.9 | 32.9 | 14.9 | 12.0 |
| LINE (%) | 58.2 | 62.9 | 27.9 | 11.6 | 11.8 |
| relic LINE (%) | 27.8 | 48.5 | 22.4 | 9.1 | 9.0 |
| LTR (%) | 73.0 | 70.0 | 36.5 | 26.9 | 18.0 |
| relic LTR (%) | 55.0 | 58.9 | 28.1 | 16.1 | 14.0 |
| Unknown (%) | 48.7 | 42.1 | 29.5 | 13.7 | 10.9 |

**Table S10. Expanded, contracted, other gene counts in five species (from CAFÉ analysis) in high or low synteny regions.** The genes that overlap in two regions were not included.

|  |  | **high synteny** | **low synteny** |
| --- | --- | --- | --- |
| *M. chlorophos* | expanded genes | 1,621 | 2,852 |
|  | contracted genes | 315 | 489 |
|  | other genes | 5,124 | 3,795 |
| *M. indigotica* | expanded genes | 1,145 | 3,091 |
|  | contracted genes | 300 | 433 |
|  | other genes | 5,288 | 4,046 |
| *M. kentingensis* | expanded genes | 979 | 4,122 |
|  | contracted genes | 181 | 529 |
|  | other genes | 3,787 | 5,400 |
| *M. sanguinolenta* | expanded genes | 2,985 | 7,333 |
|  | contracted genes | 389 | 671 |
|  | other genes | 7,026 | 6,900 |
| *M. venus* | expanded genes | 4,008 | 9,001 |
|  | contracted genes | 230 | 339 |
|  | other genes | 6,875 | 6,020 |

**Table S11. Annotation of gene function in the 29 OGs containing at least one upregulated gene in the four bioluminescent *Mycena* species.**

| **OG number** | **OG annotation** |
| --- | --- |
| OG0009249 | luciferase |
| OG0000706 | FAD/NAD(P)-binding domain-containing protein |
| OG0002489 | hispidin synthase/polyketide synthetase |
| OG0000386 | FAD-binding domain-containing protein/reticuline oxidase/ Glucooligosaccharide oxidase |
| OG0000288 | FAD/NAD(P)-binding domain-containing/ halogenase |
| OG0000215 | FAD/NAD(P)-binding domain-containing protein/ monooxygenase |
| OG0001192 | NAD(P)-binding protein/D-xylose 1-dehydrogenase (NADP(+)) 2 |
| OG0001818 | Eukaryotic aspartyl protease/ acid protease |
| OG0000591 | Peptidase S41 family protein ustP |
| OG0000147 | ABC transporter transmembrane region |
| OG0000398 | transporter |
| OG0000593 | predicted protein (pfam:SNARE_assoc) |
| OG0000348 | predicted protein (only 4 gene have pfam annotation) |
| OG0000253 | predicted protein (no any species has Pfam) |
| OG0000022 | predicted protein (pfam:Fungal_trans +Zn_clus) |
| OG0000345 | GH79 |
| OG0000143 | GH67/alpha-L-rhamnosidase-like protein |
| OG0000048 | glycoside hydrolase/retinol dehydrogenase 12/delta-9 fatty acid desaturase protein |
| OG0000014 | carboxypeptidase s1/Uncharacterized hydrolase/acid protease |
| OG0000134 | cytochrome P450/ Bifunctional P-450/NADPH-P450 reductase |
| OG0000005 | cytochrome P450 |
| OG0000066 | Zinc finger protein |
| OG0000034 | transcription initiation factor IIA gamma subunit/predicted protein |
| OG0000026 | acetyl-CoA synthetase-like protein/NRPS (nonribosomal peptide synthetase)-like enzyme |
| OG0000007 | ankyrin repeat protein |
| OG0000006 | transmembrane protein |
| OG0000097 | argonaute-like protein |
| OG0000004 | kinase-like protein/GH76/argonaute-like protein |
| OG0000000 | short-chain dehydrogenase/reductase family protein/ubiquitin family protein/glycoside hydrolase family 7 protein/MFS general substrate transporter/NAD(P)-binding protein |

**Table S12. Scaffolds containing telomeric repeats at ends.**

| **Scaffold name** | **Scaffold length** | **End** | **Repeat start** | **Repeat end** | **Number of copies** |
| --- | --- | --- | --- | --- | --- |
| Mc.scaff0010 | 1,898,893 | 5’ | 1 | 189 | 30.8 |
| Mc.scaff0018 | 1,039,458 | 5’ | 1 | 160 | 26.8 |
| Mind.scaff0001 | 6,537,710 | 5’ | 10 | 117 | 18.3 |
| Mind.scaff0001 | 6,537,710 | 3’ | 6,537,600 | 6,537,710 | 19.2 |
| Mind.scaff0002 | 6,307,876 | 3’ | 6,307,754 | 6,307,876 | 21 |
| Mind.scaff0003 | 5,994,024 | 3’ | 5,993,915 | 5,994,024 | 19.3 |
| Mind.scaff0006 | 5,133,973 | 3’ | 5,133,894 | 5,133,973 | 14 |
| Mind.scaff0007 | 5,077,951 | 3’ | 5,077,842 | 5,077,951 | 18.8 |
| Mind.scaff0008 | 5,050,887 | 5’ | 1 | 97 | 17 |
| Mind.scaff0009 | 4,505,476 | 3’ | 4,505,343 | 4,505,476 | 22.5 |
| Mind.scaff0011 | 3,413,854 | 3’ | 3,413,723 | 3,413,854 | 22.3 |
| Mind.scaff0013 | 2,685,629 | 3’ | 2,685,517 | 2,685,629 | 19.8 |
| Mind.scaff0015 | 1,639,376 | 5’ | 8 | 133 | 21.3 |
| Mk.scaff0001 | 5,940,114 | 5’ | 1 | 136 | 23 |
| Mk.scaff0001 | 5,940,114 | 3’ | 5,939,956 | 5,940,114 | 26.5 |
| Mk.scaff0003 | 4,788,751 | 5’ | 1 | 165 | 27.5 |
| Mk.scaff0005 | 4,011,168 | 3’ | 4,011,018 | 4,011,168 | 25.2 |
| Mk.scaff0006 | 3,839,597 | 5’ | 1 | 144 | 23.7 |
| Mk.scaff0008 | 3,039,429 | 5’ | 1 | 159 | 26.7 |
| Mk.scaff0009 | 2,763,528 | 5’ | 1 | 165 | 27.8 |
| Mk.scaff0012 | 2,261,502 | 5’ | 1 | 159 | 26.7 |
| Mk.scaff0013 | 2,164,973 | 3’ | 2,164,823 | 2,164,973 | 25.2 |
| Mk.scaff0014 | 2,112,720 | 3’ | 2,112,566 | 2,112,720 | 26.3 |
| Mk.scaff0016 | 1,974,397 | 5’ | 2 | 156 | 26.3 |
| Mk.scaff0017 | 1,721,327 | 3’ | 1,721,151 | 1,721,327 | 29.5 |
| Mk.scaff0019 | 1,349,653 | 5’ | 1 | 151 | 25 |
| Msan.scaff0001 | 17,789,092 | 3’ | 17,789,025 | 17,789,090 | 11 |
| Msan.scaff0025 | 1,760,574 | 3’ | 1,760,504 | 1,760,574 | 11.5 |
| Msan.scaff0027 | 1,650,193 | 3’ | 1,650,124 | 1,650,193 | 11.3 |
| Mven.scaff0003 | 11,118,782 | 3’ | 11,118,677 | 11,118,782 | 17.2 |
| Mven.scaff0004 | 8,657,829 | 3’ | 8,657,762 | 8,657,829 | 11.3 |
| Mven.scaff0007 | 6,670,972 | 5’ | 1 | 70 | 11.8 |
| Mven.scaff0008 | 6,224,985 | 3’ | 6,224,891 | 6,224,985 | 15.8 |
| Mven.scaff0011 | 5,452,461 | 3’ | 5,452,365 | 5,452,461 | 16 |
| Mven.scaff0012 | 5,262,452 | 5’ | 1 | 125 | 20.7 |
| Mven.scaff0013 | 4,843,808 | 3’ | 4,843,711 | 4,843,808 | 16.3 |
| Mven.scaff0014 | 4,029,179 | 5’ | 1 | 100 | 16.7 |
| Mven.scaff0015 | 3,840,008 | 3’ | 3,839,974 | 3,840,008 | 5.8 |
| Mven.scaff0020 | 3,032,789 | 3’ | 3,032,670 | 3,032,789 | 20 |
| Mven.scaff0028 | 2,074,661 | 5’ | 1 | 94 | 15.7 |

**Table S13. Bisulfite conversion rate from bisulfite sequences aligned to the genome assemblies of *M.* *kentingensis*.**

| **Species** | **Replicate** | **Number of raw reads** | **After PCR duplicate removing and adapter trimming** | **Number of uniquely mapped** | **Mappability (%)** | **DNA methylation level (%)** | | | **Coverage (depth) per strand (X)** | **Bisulfite conversion rate (%)*** |
| --- | --- | --- | --- | --- | --- | --- | --- | --- | --- | --- |
|  |  |  |  |  |  | **CG** | **CHG** | **CHH** |  |  |
| *M. kentingensis* | 1 | 39,272,420 | 35,864,994 | 21,411,409 | 59.7 | 15 | 1.7 | 1.75 | 19.23 | 98.95 |
|  | 2 | 37,212,556 | 33,627,150 | 20,205,636 | 60.09 | 14.48 | 1.6 | 1.62 | 18.02 | 99.06 |

*The bisulfite conversion rate was estimated by spiking in unmethylated Lambda Phage DNA into the *Mycena* BS libraries. We mapped BS-reads to Lambda Phage genome, so any unconverted cytosines revealed the failed bisulfite conversion and were considered false positives.

**Dataset S1 (separate file). Statistics of RNAseq reads and coverage.**

**Dataset S2 (separate file). OG number in 42 species.**

**Dataset S3 (separate file). Differentially-expressed genes between mycelia with different bioluminescent intensities in four *Mycena* species**. For *M. kentingensis* and *M. chlorophos,* genes were identified between two tissues with contrasting bioluminescence intensity*.* For *M. sanguinolenta* and *M. venus,* genes were identified by significant correlation between bioluminescent intensity and expression level.

**Dataset S4 (separate file). Differentially-expressed genes between the cap and stipe of the fruit body in *Mycena* *kentingensis*.** The genes with red color located in luciferase cluster.

**Dataset S5 (separate file). The genes assigned to module50 from the analysis of WGCNA during the developmental stages in *M. kentingensis*.** The genes with red color located in luciferase cluster.

**Dataset S6 (separate file). Molecular function of GO terms enriched in proteins of 589 OGs expanded at the origin of the mycenoid lineage.**

**Dataset S7 (separate file). Pfam domain number in 42 species.**

**Dataset S8 (separate file).** **537 protein domains enriched in the mycenoid lineage**. Protein domains expanded (Wilcoxon rank sum test, p < 0.01; 1-fold higher copy number) in six mycenoid species compared to the other 36 species.

**Dataset S9 (separate file). The diameter of the mycelium and its bioluminescence for seven days.**
